# A mechanistic view of collective filament motion in active nematic networks

**DOI:** 10.1101/732909

**Authors:** Moritz Striebel, Isabella R. Graf, Erwin Frey

## Abstract

Protein filament networks are structures crucial for force generation and cell shape. A central open question is how collective filament dynamics emerges from interactions between individual network constituents. To address this question we study a minimal but generic model for a nematic network where filament sliding is driven by the action of motor proteins. Our theoretical analysis shows how the interplay between viscous drag on filaments and motor-induced forces governs force propagation through such interconnected filament networks. We find that the ratio between these antagonistic forces establishes the range of filament interaction, which determines how the local filament velocity depends on the polarity of the surrounding network. This force propagation mechanism implies that the polarity-independent sliding observed in Xenopus egg extracts, and in-vitro experiments with purified components, is a consequence of a large force propagation length. We suggest how our predictions can be tested by tangible *in vitro* experiments whose feasibility is assessed with the help of simulations and an accompanying theoretical analysis.

## 1 Introduction

Living cells have the remarkable ability to actively change their shape, and to generate forces and motion. A key component enabling cells to exhibit these stunning mechanical properties is the cytoskeleton. This structure is built out of various proteins and forms diverse functional networks consisting of polymer filaments such as actin and microtubules, motor proteins, and associated proteins ^1, 2^. The motor proteins expend chemical energy to generate forces that act on the cytosceletal filaments ^3–5^. In particular, motors that have two binding domains, e.g. kinesin-5, can walk along two filaments at once, causing filaments of opposite polarity to slide past one another ^6^.

To understand the non-equilibrium physics underlying the dynamics of motor–filament systems, it has proven fruitful to study reconstituted systems of purified components *in vitro* ^7–10^. Despite their reduced complexity, these systems still self-organize into intricate patterns and structures reminiscent of those found in living cells. But how is their collective behavior at the macroscopic level linked to the interactions between individual filaments and motors? What are the underlying mechanisms? To provide an answer, we focus here on a generic class of systems in which filaments exhibit nematic order and motors drive relative sliding of filaments. A prominent representative of this class is the poleward flux of microtubules in Xenopus mitotic spindles ^11–13^. This process has been attributed to antiparallel, motor-driven interactions between filaments, especially if the motor protein dynein is inhibited ^11, 14, 15^. A quite puzzling observation made in these systems was the correlation — or rather, the lack of correlation — between filament speed and network polarity, i.e. the ratio of parallel to antiparallel filaments. Although filament motion is induced by sliding antiparallel filaments past each other, polarity was observed to have barely any influence on the filament speed ^14, 16, 17^. This surprising behavior was recently replicated in a system of purified components composed of the kinesin-14 XCTK2 and microtubules, and interpreted in terms of a hydrodynamic theory for heavily crosslinked filament networks ^18^. These observations are at variance with previous predictions for dilute filament networks, where filament motion depends linearly on the local polarity ^19–21^. How can these conflicting results be reconciled? What are the biophysical mechanisms determining the relation between filament speed and network polarity?

To gain insight into these important questions we study a minimal but generic model consisting of nematically ordered cytoskeletal filaments (like microtubules) and molecular motors (like kinesin-5) that are capable of crosslinking and sliding antiparallel filaments apart. Our mathematical analysis of this theoretical model shows that the interplay between motor-induced forces and viscous drag acting on the filaments determines the relation between filament velocity and the polarity of filaments. Depending on the relative strengths of these forces, we find that the velocity–polarity relation varies continuously between a local and a global law. Our theory reveals the mechanism that underlies this relation between filament velocity and network polarity: For high motor-induced forces and small fluid drag, local forces on the filaments propagate through the strongly interconnected network without dissipation and thereby influence the overall network dynamics. In contrast, for small motor–induced forces or high fluid drag, local forces are quickly damped and only influence the local dynamics. This mechanism provides a deeper understanding of the link between collective filament dynamics and molecular interactions. Moreover, it reconciles previously conflicting results for the velocity-polarity relation in the limit of dilute ^19, 20, 20^ and heavily crosslinked systems ^14, 18, 22^. Strikingly, our theoretical analysis shows that the insensitivity of filament velocities to changes in the network polarity, which was reported for the spindle ^14, 16^ and *in vitro* systems ^18^, occurs in a biologically relevant parameter range. In addition our theory predicts how the ratio between the spectrum of measured polarities and filament speeds depends on the ratio of drag to motor-induced forces in the system. We suggest an *in-vitro* experiment to validate those predictions. The feasibility of this experiment is assessed with the help of computer simulations and an accompanying theory.

## 2 Results

### Biophysical agent-based model of motor-induced filament movement

We are interested in understanding how the interplay between viscous drag and molecular forces between cytoskeletal filaments, mediated by molecular motors, drives the internal dynamics of filament networks. Specifically, we focus on reconstituted *in vitro* systems consisting of microtubules and motors capable of crosslinking neighboring filaments and sliding them apart, c.f. Fig. 1 A, B. Such motor proteins can walk on both filaments simultaneously, so that the forces generated between filaments depend on their relative orientation (Fig. 1 A). *In vitro* such microtubule-motor mixtures were observed to self-organize into a nematic network, where neighboring filaments may be disposed approximately parallel or antiparallel ^18^.

**Figure 1:**
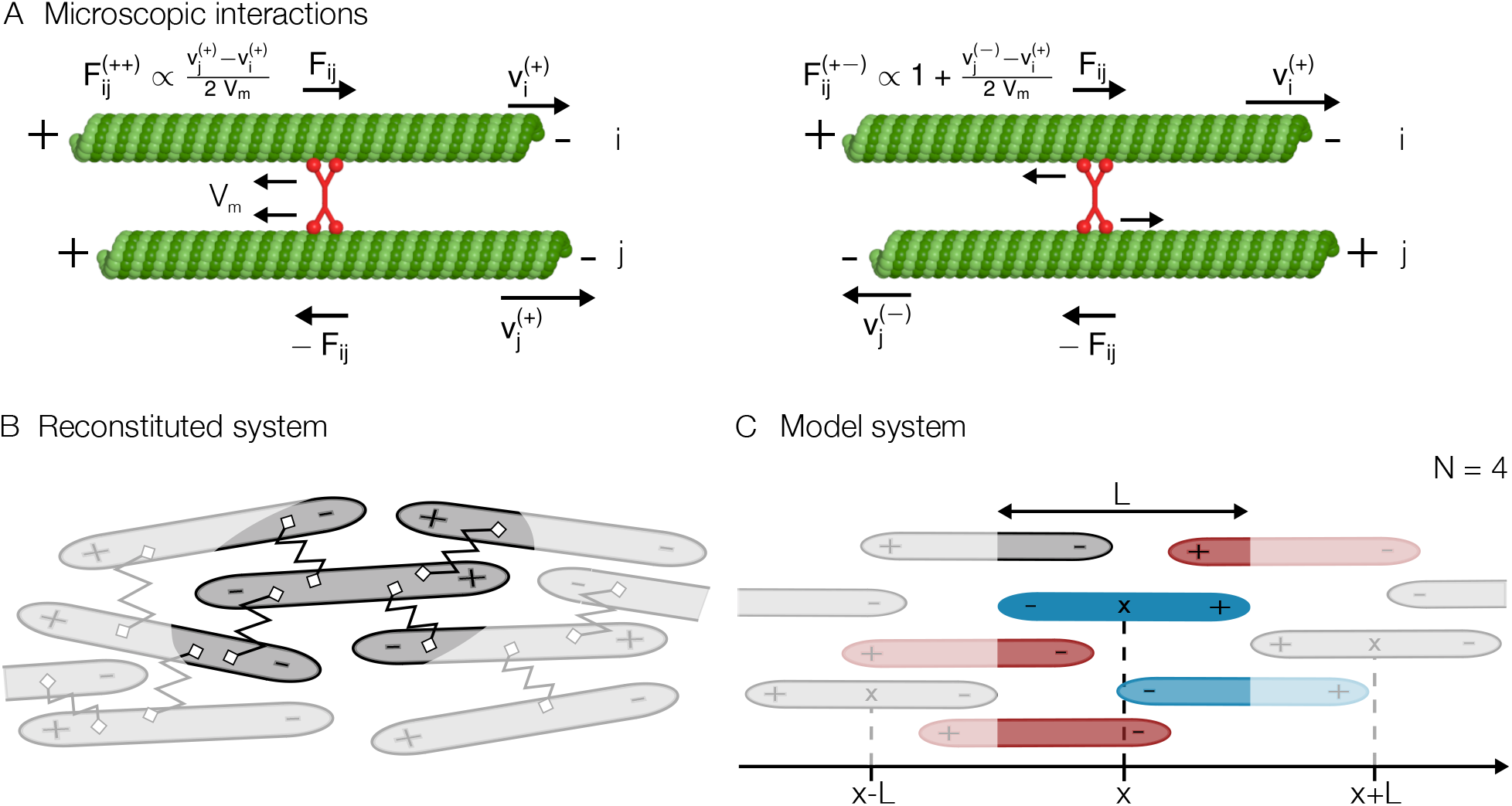
Biophysical model for motor-driven filament motion. (**A**) Microscopic, motor-mediated interactions between microtubules. Neighboring microtubules are connected by motors (red) which walk towards the microtubule’s (green) plus end with velocity *V*_m_. A motor exerts zero force if filament motion is such that the motor is not stretched. (**A -left**) A motor connecting two parallel microtubules counteracts relative motion between the filaments. (**A -right**) In contrast, two antiparallel microtubules connected by a motor are slid apart. The force falls to zero once their relative velocity equals twice the motor velocity (−2 *V*_m_). (**B**) Sketch of a microtubule-motor mixture in a nematically aligned state. The springs denote motors crosslinking neighboring filaments. The highlighted region includes all interactions of the center microtubule. (**C**) The one-dimensional model system. Possible interaction partners of the microtubule in the center (dark blue) are in the highlighted region. To account for the reduced number of interaction partners in the experimental filament network, we draw on average *N* out of all possible partners to interact (interaction partners are highlighted in color, parallel interaction partners in light blue and antiparallel interaction partners in light red).

Motivated by the nematic order of these filament networks, we set up a biophysical agent-based model, which is effectively one-dimensional. We consider a system of size *S* where the filaments (microtubules) are assumed to be rigid polar rods of fixed length *L*, oriented with their plus end either to the left (+) or right (−); see Fig. 1 C. Hence, the dynamics of each polar filament *i* is determined (solely) by its velocity 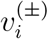. Relative motion between filaments is caused by molecular motors that walk on these crosslinked filaments and thereby exert forces. In-vitro assays involving pairs of isolated microtubules cross-linked by kinesin-5 motors reveal that: (a) Kinesin-5 has the ability to walk simultaneously on both microtubules with approximately the zero-load velocity *V*_m_, (b) antiparallel microtubules are pushed apart with a relative velocity of ~ 2*V*_m_, and (c) parallel microtubules remain static ^6^. Integrating this information with experiments showing a linear force-velocity relation for kinesin motors ^23–26^, we assume that the forces between two crosslinked parallel (±±) and antiparallel (±∓) filaments per motor are given by

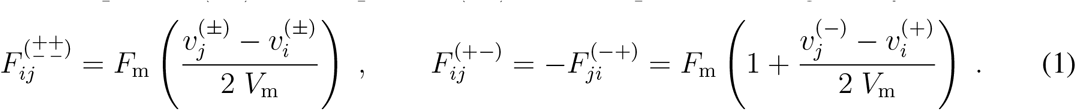

Here *F*_*ij*_ denotes the force that filament *j* exerts on filament *i*, with *F*_m_ signifying the motor stall force; due to force balance *F*_*ij*_ = −*F*_*ji*_. These forces vanish if the relative motion of the filaments does not induce strain in the crosslinking motors. While for parallel filaments this is the case if the filaments move at the same speed, a motor walking on antiparallel filaments is not strained if these slide apart with relative velocity 2*V*_m_, i.e. 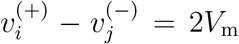. On the other hand, the maximal force between two filaments corresponds to the stall force, *F*_m_, which is defined as the force between two antiparallel filaments fixed at their relative position 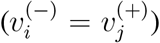. In that case the motor heads move apart until the motor stalls and exerts its maximal force on the filaments. An analogous situation occurs if a motor is attached to two parallel filaments which move with a relative speed 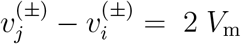. So the corresponding force is also *F*_m_.

The velocity 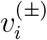 of a specific microtubule *i* in the network is determined by the force balance equation

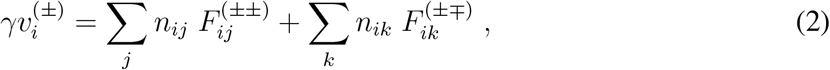

where *γ* denotes the fluid drag coefficient and *n*_*ij*_ the number of motors crosslinking microtubule *i* and *j*. The sums run over all parallel and antiparallel interaction partners of microtubule *i*, respectively. In general, the number of interaction partners as well as the strength of their interaction can depend on a variety of factors. For example, the interactions are influenced by the density of motors in the cytosolic volume as well as along the filament, or the local structure of the filament network. Inclusion of all these factors would lead to a microscopic description with many unknown parameters. Focusing on the mechanistic basis of filament motion here, we make the following two assumptions (c.f. Fig. 1 C): First, we consider a homogeneous motor density in the cytosolic volume and along the microtubules. Thus, we describe motors effectively by a constant density, with on average *N*_*m*_ motors per filament. Second, we assume that all filaments have on average *N* interaction partners that are drawn randomly. This on average accounts for the limited number of neighbors in the three-dimensional network structure.

To gain initial insight into the dynamics of microtubules, we simplify the system even further using a local, continuum mean-field approximation that neglects any lateral displacement between crosslinked filaments. In the continuum description, each microtubule *i* is identified by its midpoint position *x*_*i*_. As a crude simplification, we assume that all crosslinked, equally oriented filaments passing through position *x* move at (roughly) the same velocity *v*^(±)^(*x*). This entails that the forces between all parallel filaments, *F*^(±±)^(*x*), vanish. Denoting the fraction of filaments at position *x* oriented in (±) direction by *φ*^(±)^(*x*), Eq. 2 then simplifies to *γv*^(+)^(*x*) = *N*_m_*φ*^(−)^(*x*)*F*^(+−)^(*x*) and *γv*^−^(*x*) = *N*_m_*φ*^(+)^(*x*)*F*^(−+)^(*x*) with *N*_m_ denoting the number of motors per filament as above. Inserting the force velocity equation, Eq. 1, and solving for the velocity yields *v*^(+)^(*x*) ∝ 1 − *P*(*x*) and *v*^(−)^(*x*) ∝ 1 + *P*(*x*), where we defined *P*(*x*) = *φ*^(+)^(*x*) − *φ*^(−)^(*x*) as the local network polarity at position *x*. Hence, the central result of this local mean-field analysis, which will ultimately turn out to be oversimplified, is a linear dependence of the local velocities on the local polarity. This result corresponds to the intuition that forces between filaments — and their relative motions — strongly depends on their relative orientation. In particular, while antiparallel interactions between two filaments introduce motion of both filaments, parallel filaments remain static. As a consequence, filaments with a higher number of antiparallel interactions are expected to exhibit an enhanced speed. However, as we will see next, this intuition is in conflict with numerical simulations (in the biologically relevant parameter regime) as well as with experimental findings for heavily crosslinked filament gels ^18^.

### The agent-based model can describe the weak velocity-polarity sensitivity

To test whether our model is capable of describing the observations in heavily crosslinked filament networks, we solved the full set of coupled linear equations (Eq. 2) for a one-dimensional network numerically. In order to compare our results to experimental data, we assessed the model parameters as follows: First, we determined the mean number of interaction partners per filament. The typical maximal distance between two microtubules connected by a sliding motor is estimated to be of the order of the tail length of kinesin-5, ~ 0.1μm, ^5^ plus two times the microtubule radius, ~ 0.024 μm, ^27^. Together with the typical microtubule length, estimated to be ~ 6 − 7 μm, these values yield an interaction volume of approximately 1/3 μm^3^. Fürthauer and collaborators argue that the number density of filaments in their experimental setup is approximately 17/μm^3^ ^18^. So, all in all, we estimate that there are *N* ≈ 5.5 interaction partners per filament. In an analogous manner, we assessed the number of microtubules in our one-dimensional representation of the experimental chamber of length 400 μm to be ~ 400. Those filaments are placed randomly as described below (section “*In silico* experiment: Random polarity field”) and experience a drag coefficient of *γ* = 0.5 pN s/μm ^28–30^. As motor parameters we use *V*_m_ = 20 nm/s ^6^, *F*_m_ ~ 1 pN ^5^ and *N*_m_ = 25 as the average number of motors per filament ^18^.

Using these parameters we performed numerical simulations, and found good agreement with experimental results (compare Fig. 2 and Fig. 2 in Ref. ^18^). In particular, the average filament speed (filled black circles in Fig. 2) is found to be independent of the local polarity. This clearly contradicts the local mean-field theory as discussed above (see section “Biophysical agent-based model of motor-induced filament movement”). To assess why this simplified local view is misleading, we next give a comprehensive mathematical analysis of the full agent-based model.

**Figure 2:**
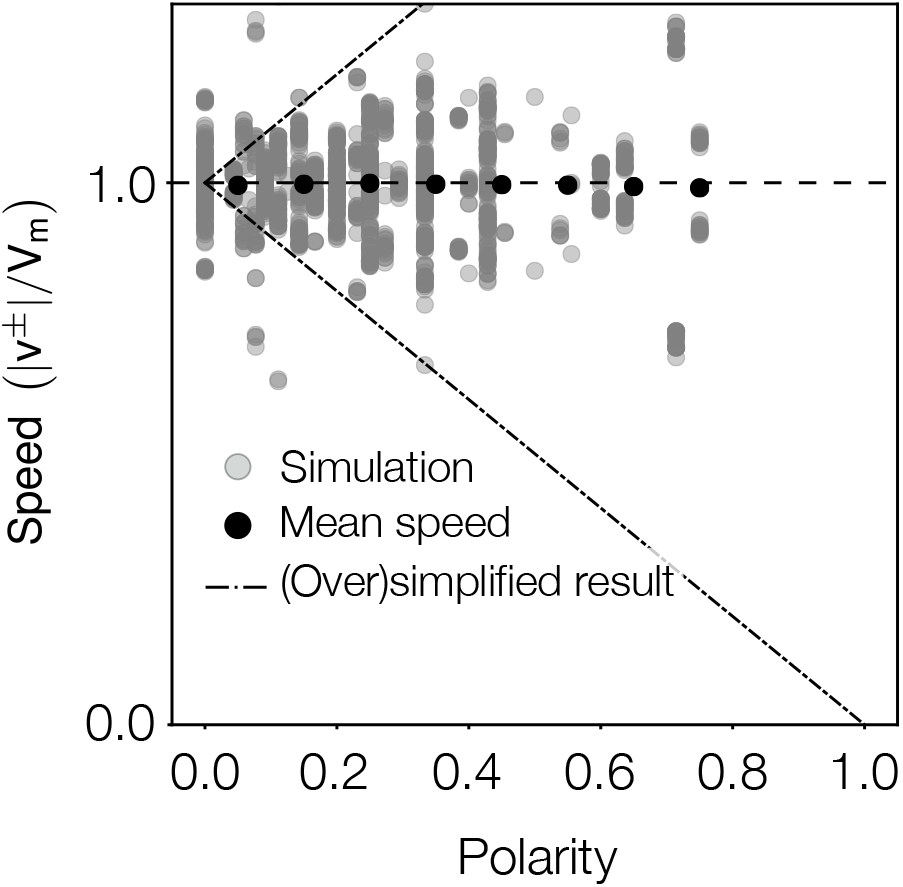
Local microtubule speed vs. local polarity obtained by numerically solving the full set of coupled linear equations (Eq. 2) for a one-dimensional microtubule network. The microtubule network is generated as described in section “*In silico* experiment: Random polarity field”. Gray dots represent individual measurements, black dots show the average speed binned for local polarities (bin size Δ*P* = 0.1). In contrast to the oversimplified discussion (dashed-dotted lines), the velocity does not depend linearly on the local polarity. Instead, the average speed is mostly independent of the local polarity. Note that the vertical stripes are artefacts arising from the discrete nature of the agent-based simulation: Due to the finite number of filaments in an interval [*x, x* + Δ*x*] the polarity can only take on discrete values.

### Non-local continuum theory

It is evident that in the simplified local mean-field analysis discussed above we neglected the finite extension of filaments. Actually, two filaments which pass through the same location do not necessarily have the same midpoint position. While they share some overlap, they will interact with different neighbors at different positions. If all filaments have the same length *L*, a filament with midpoint at position *x* can interact with filaments whose mid-points lie in the interval [*x* − *L*, *x* + *L*] (cf. Fig 1 C). In this way, the velocities of filaments located at different spatial positions are coupled, leading to non-local correlation effects that could explain the weak dependence of filament speed on local polarity.

Motivated by this heuristic argument, we set out to formulate a continuum theory that quantifies the non-local coupling between the filament velocities (*v*^±^(*x*)) and densities (*ρ*^(±)^(*x*)). To this end, we rewrote the local balance equation, Eq. 2, assuming a continuum limit.

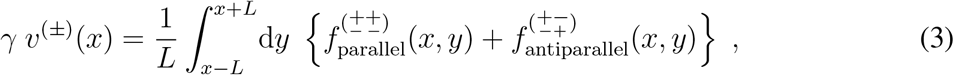

where the local forces are given by

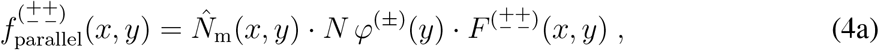

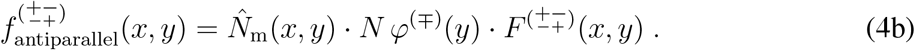

Here, the force a motor exerts on the filaments it crosslinks is simply given by the continuum version of Eq. 1, e.g. *F*^(±±)^(*x*, *y*) = *F*_m_ [*v*^(±)^(*y*) − *v*^(±)^(*x*)]/(2 *V*_m_). The second factor in Eq. 4 accounts for the expected number of interaction partners at position *y*, given by the number fraction of filaments with the respective polarity *φ*^(±)^(*y*) multiplied by the average number *N* of interaction partners: *N φ*^(±)^(*y*). For this functional form to apply, we implicitly assumed that the filament network is not sparse, i.e., that there is always a sufficient number of interaction partners, namely more than *N*, available. The number fraction can be written in terms of the filament densities as *φ*^(±)^ = *ρ*^(±)^/(*ρ*^(+)^ + *ρ*^(−)^). The first factor in Eq. 4, 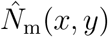, specifies the average number of motor proteins mediating the interaction between a pair of filaments located at positions *x* and *y*. This number is determined by the size of the overlapping region, *L*_ov_ = *L*−|*x*−*y*|, and the number of motors per filament, *N*_m_. Since all the available motors on a filament have to be shared among all of its *N* interaction partners, only *N*_m_*/N* are available for the interaction with any specific filament. Hence, assuming a uniform motor distribution along each microtubule, the effective number of motors crosslinking a filament pair is on average given by 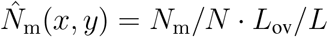.

Based on this non-local continuum representation of our agent-based model, we seek a quantitative understanding of how the opposing forces in the filament network give rise to collective (uniform) motion. Ultimately, our goal is to provide an explicit expression relating the polarity and velocity fields.

### Analytic solution for motor-induced filament movement

In this section, we present an analytic solution to our non-local continuum description (Eq. 3). We restrict our analysis to the limit where the system size is large compared to the filament length *L* and to all other intrinsic length scales of the system we might encounter in the course of the mathematical analysis. Making use of complex calculus, in this limit it is possible to find an explicit expression for the velocity field *v*^(±)^(*x*) in terms of the polarity field *P*(*x*). This expression, thus, constitutes a velocity-polarity relation which quantifies how the polarity field affects the velocities.

In an experimentally reasonable parameter regime one finds an approximate expression which reads (for a detailed analysis see “An analytic solution for filament motion in a nematic network)”:

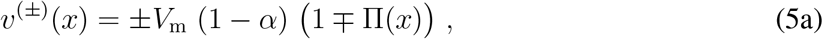

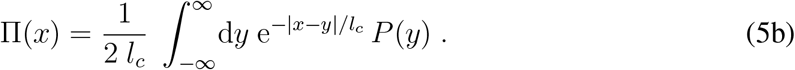

where 1/*α* := 1 + 12 (*l*_*c*_/*L*)^2^; for biologically plausible parameter values one has *α* ≪ 1. Importantly, Eq. 5a shows that the motion of filaments is neither solely dependent on the local polarity nor fully independent of the polarity field. Instead, the local filament velocities, *v*^(±)^(*x*), now depend in a non-local way on the polarity, *P*(*y*), as specified by the convolution integral (weighted average), Π(*x*), with an exponential kernel 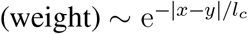. To emphasize this non-local dependence of the velocities on the polarity, we refer to Π(*x*) as the *ambient polarity* in the following. The characteristic interaction range *l*_*c*_, over which the polarity field is averaged, is given by

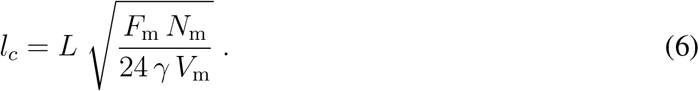

It is set by the ratio of the total average force exerted by motors on a microtubule, *F*_m_ *N*_m_, to the drag imposed on the microtubule by the surrounding fluid, *γ V*_m_. Furthermore, it can be interpreted as the length scale over which motion generated by antiparallel filament sliding is propagated by parallel filament interactions through the network. As a result, the interaction range *l*_*c*_ reflects the antagonism between the combined effect of motion-generating forces (antiparallel interactions) and motion-propagating forces (parallel interactions), and the attenuation of force propagation in the filament network mediated by viscous drag. This antagonism is captured by the spatial average of the polarity field which effectively corresponds to a low-pass filter. Due to averaging over local polarities, high-frequency fluctuations in the spatial polarity profile are filtered out and, hence, do not contribute to the velocity. Explicitly, by Fourier transforming Eq. 5b we find a Lorentzian Fourier weight

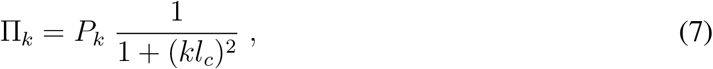

where *k* denotes the wave number. Hence, the characteristic frequency of the low-pass filter is proportional to the reciprocal of the characteristic length, 1/*l*_*c*_, implying that the larger *l*_*c*_ the stronger the filter and the less relevant local fluctuations in the polarity. To put it another way, the speed of a filament at position *x* depends only on the local “view” of the polarity field within a range defined by *l*_*c*_ (Fig. 3).

**Figure 3:**
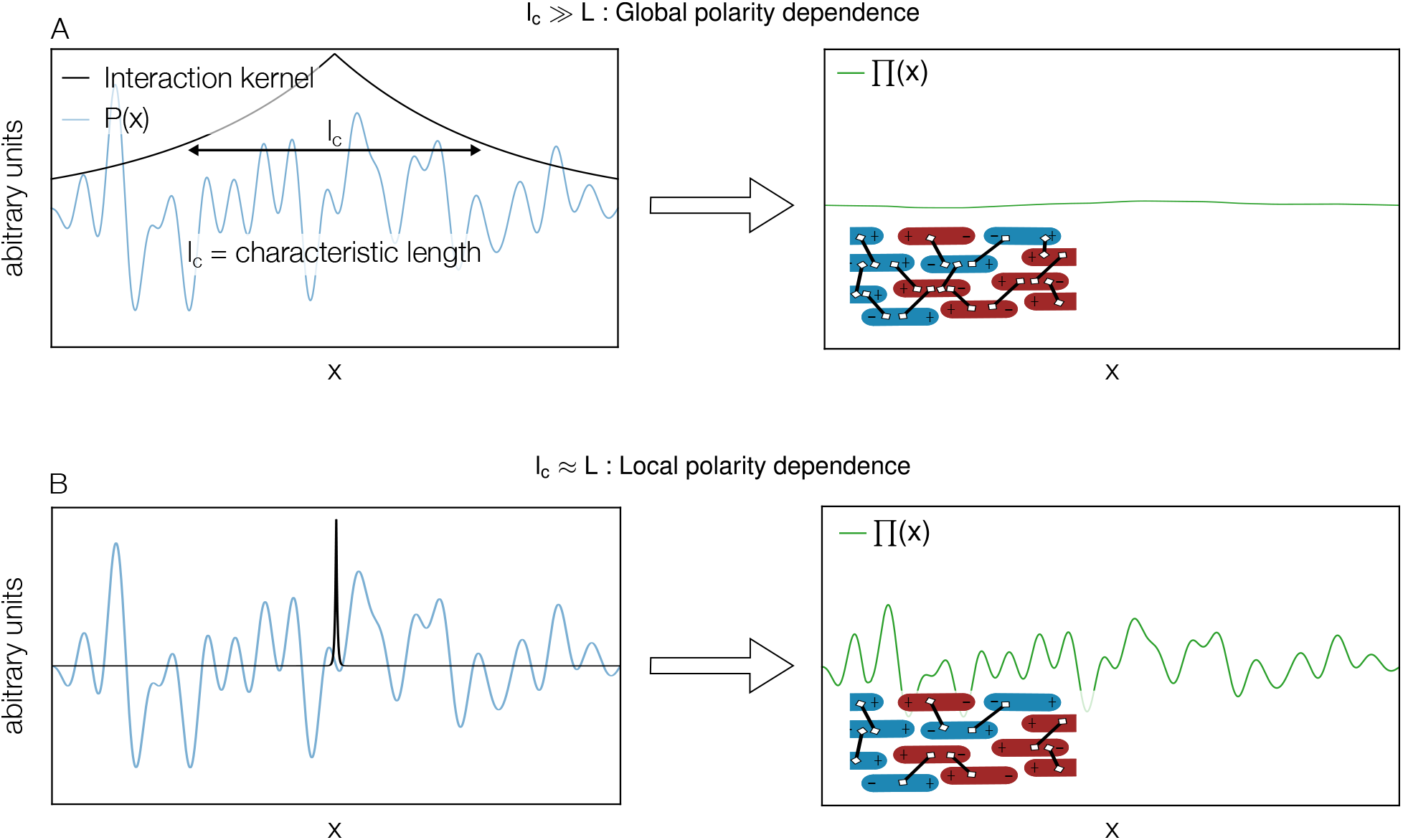
Typical polarity field, *P*(*x*), and two choices of interaction kernel, exp(−|*x* − *y*|/*l*_*c*_), characterizing global and local polarity dependence, respectively. Filaments positioned in a range of *l*_*c*_ around *x* contribute to the motion of microtubules at *x*. Depending on the ratio of average motor force exerted on a microtubule to attenuation (drag of microtubules in the fluid), the characteristic propagation length *l*_*c*_ takes different values. (**A**) For large *l*_*c*_ and polarity fields that vary randomly on length scales smaller than *l*_*c*_, this averaging yields a roughly constant ambient polarity profile, Π (*x*), and hence a roughly constant velocity p rofile. On a microscopic level, this corresponds to a heavily crosslinked filament network (inset). (**B**) In the limit of small *l*_*c*_ only the local environment, i.e., the direct interaction partners, has an influence on the microtubule motion. The ambient polarity field (velocity field) varies as the polarity field varies.

To gain an impression of how the interplay between the different forces in the network leads to the non-local effects, it is helpful to consider the limiting cases of large and small *l*_*c*_, respectively. For large *l*_*c*_, motor forces dominate viscous drag (*F*_m_*N*_m_ ≫ *γV*_m_). Then, either due to weak dissipation or strong motor-mediated filament coupling, parallel crosslinked filaments translate the motion, generated by interactions between antiparallel filaments, over long distances (~ *l*_*c*_). As a result, motion generated at one position in the network propagates through the entire network. In the asymptotic limit *l*_*c*_ → ∞, the velocity-polarity relation (Eq. 5a) reduces to *v*^(±)^ = *±V*_*m*_^1^, confirming recently published findings for a heavily crosslinked network ^18^. In contrast, for small *l*_*c*_ (*F*_m_*N*_m_ ≪ *γV*_m_) force generated at a certain position in the network has only a local effect. Forces generated by antiparallel interactions cannot propagate through the network either due to strong dissipation or a lack of parallel filament interactions. In this limit, the velocity-polarity relation reduces to the result obtained with the local mean-field theory discussed in “Biophysical agent-based model of motor-induced filament movement”. This relation agrees with the velocity-polarity relation found for dilute filament networks where only local bundles of filaments are considered ^20, 21 2^.

### Interpretation of the velocity-polarity relation

With regard to previous results, our considerations offer a solution to the seemingly contradictory behavior of dilute and heavily crosslinked networks. More specifically, our results identify a common mechanism for collective filament dynamics: Due to the finite extension of the microtubules, one microtubule can be crosslinked with several others whose center positions are spread over a region up to twice the microtubule length (Fig. 1C). As a result, although microtubules at different positions might in fact not be directly linked by a motor, an interaction between them can be mediated by successive crosslinks through a chain of microtubules. In this way, the velocity of microtubules at one position influences the velocity at a different position and information on the local polarities propagates through the system. How far this information propagates (*l*_*c*_) depends on how “effectively” movement at one position is translated into movement at a different position. The greater the efficiency, the smaller the ratio between the passive drag on microtubules in the fluid (and thus the attenuation) and the average maximal active force exerted on one microtubule by all motors linking it to other microtubules.

Taken together, our results shed light on the question of what determines the local speed of microtubules in a nematic network: Generally, it is neither the local polarity, *P*(*x*), that determines the velocity of microtubules at a certain position nor the overall polarity in the system, *P*_glob_. Instead the ambient polarity, Π(*x*), is informative. The ambient polarity corresponds to an average of the polarity with a weight that decays exponentially with the distance from the position of interest (see Eq. 5b). The characteristic decay length, *l*_*c*_, is proportional to the filament length *L*, and increases with the ratio of the motor-force on a microtubule, *F*_*m*_*N*_*m*_, to the fluid drag, *γV*_*m*_. In general, for a finite decay length and a spatially varying polarity profile, the ambient polarity also varies in space. As can be inferred from Eq. 5b, for larger values of *l*_*c*_, a larger region of space contributes to the ambient polarity (see also Fig. 3). Accordingly, the ambient polarity then corresponds to an average of the local polarity over more positions. As a result, for a fixed spatial polarity profile, the range of values of the ambient polarity decreases with increasing characteristic propagation length *l*_*c*_. Due to the linear relationship between the velocities and the ambient polarity, Eq. 5a, the same holds true for the range of velocities.

In the following, we illustrate these predictions with the help of two examples. First, we consider a spatially linear polarity profile. Besides being an instructive case, this polarity profile is of biological relevance. It resembles the measured, approximately linear polarity profiles in the mitotic spindle (see section “Summary and discussion”). As a complement, the setup of the second example is designed to mimic typical *in vitro* experiments. In order to make testable predictions we analyze the suggested (idealized) experiment in detail and focus on quantities which we believe to be accessible in experiments.

### A simple example: The linear polarity profile

Our theory predicts that the range of velocities decreases with increasing characteristic propagation length, *l*_*c*_. To demonstrate this correlation, we consider a linear polarity profile *P*(*x*) = *a* (*x − S/*2) in a finite interval *x ∈* [0, *S*] (for details see Appendix “Linear polarity profile”). As motivated above, we describe the local polarity profile in terms of its Fourier coefficients 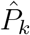. The wave numbers are now discrete, *k* ∈ ℕ, as the system is finite. The Fourier coefficients of the ambient polarity, 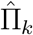, are given by the Fourier coefficient of the local polarity, 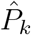, times a *k*-dependent weighting factor: 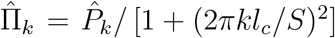 (see Appendix “Fourier representation of the ambient polarity”). Correspondingly, the ratio between the range of the local polarity 2*P*_max_ = *aS* and the range of the ambient polarity 2Π_max_ can be approximated as (see Appendix “Linear polarity profile”)

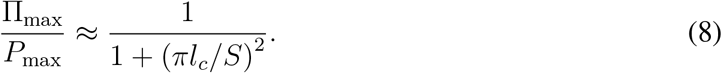

This finding confirms the intuitive expectation that with increasing characteristic length *l*_*c*_ the ambient polarity range 2Π_max_ (or analogously the velocity range 2Π_max_*V*_*m*_(1 − *α*)), should decrease relative to the local polarity range, 2*P*_max_, and the spatial profile gets “squeezed” (Fig. 4). From the approximate expression, Eq. 8, we infer that for characteristic lengths of the same order as the system size, 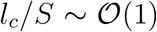, the range of the ambient polarity 2Π_max_ is only a tenth of the range of the local polarity 2*P*_max_. Due to the linear relationship between the velocity and ambient polarity, Eq. 5a, this small range of ambient polarities implies that also the velocity range for equally oriented microtubules is small. As a result, for *l*_*c*_ ≥ *S*, all equally oriented microtubules move as a collective with approximately uniform velocity. In particular, there is also movement in regions where locally the polarity is *P*(*x*) = ±1, corresponding to stretches populated only by parallel microtubules.

**Figure 4:**
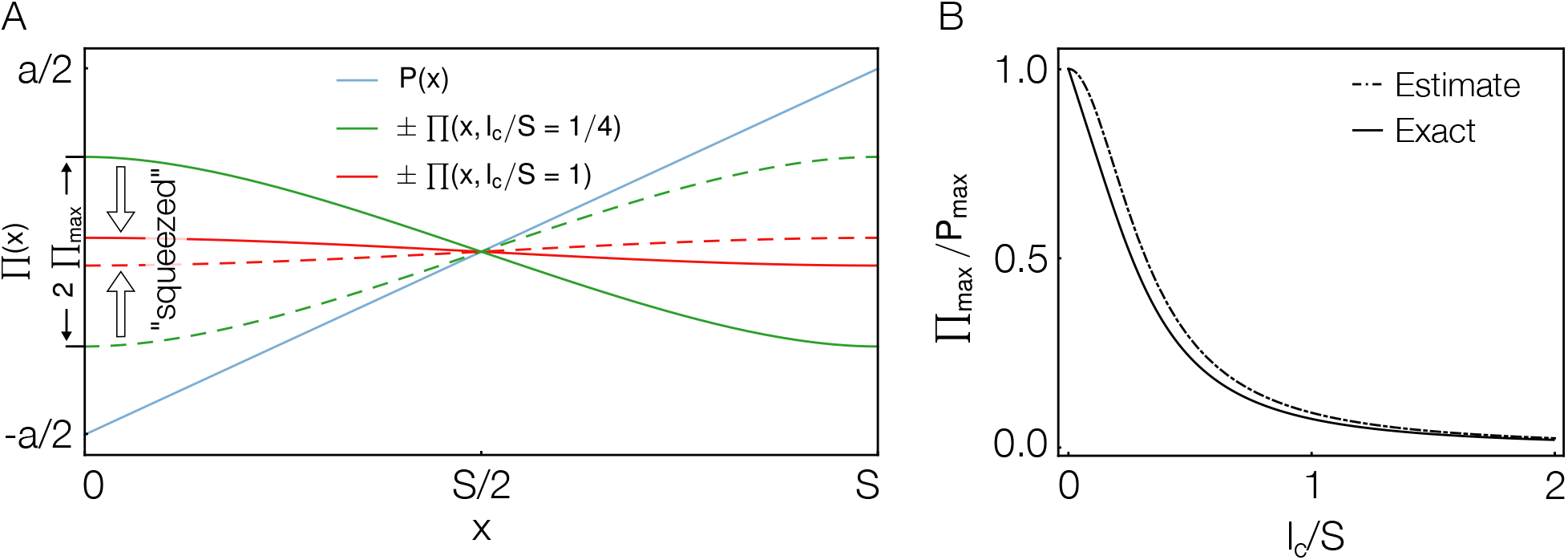
“Squeezing” of the ambient polarity in a finite system with reflecting boundary condi-tions. (**A**) Sketch of the linear spatial polarity profile, *P*(*x*) = *a* (*x−S/*2), *x* ∈ [0, *S*], together with the ambient polarity profile, Π(*x*, *l*_*c*_), for two different values of the characteristic length *l*_*c*_/*S* (normalized by the system size *S*). The solid and dashed lines indicate the solutions relevant for (+) and (−) filaments, respectively. For larger *l*_*c*_, the range of the ambient polarity, 2Π_max_, becomes more restricted. (**B**) Ratio between the range of the ambient polarity 2Π_max_ and the range of the local polarity 2*P*_max_ = *aS* plotted against *l*_*c*_/*S*. The curve is well approximated by a Lorentzian decay 1/(1 + (*πl*_*c*_/*S*)^2^) (estimate). For the exact expression, please refer to the Appendix “Linear polarity profile”. For larger *l*_*c*_/*S*, the range of the ambient polarity relative to that of the local polarity falls off rapidly.

For *in vitro* experiments with filament gels or reconstituted systems, it might not be feasible to get information on the entire spatial polarity and velocity fields. Instead, in typical experiments the local polarity and velocity are recorded only at single points in the filament gel ^17, 18, 31^. Data obtained in this way is similar to that shown in Fig. 2 where one data point corresponds to a polarity-velocity pair measured at one location in the gel. In the next section, we thus perform an *in silico* experiment where we make single velocity and polarity measurements only and do not measure the entire spatial fields. Nevertheless, the key idea motivating the setup of the *in silico* experiment is the expectation that the spectrum of measured velocities is squeezed compared to the spectrum of local polarities: Due to the filtering of short-wavelength modes, extreme values of the local polarity are averaged out and the velocity profile is smoother than the local polarity profile. In the following, we thus focus on deriving a relation between the measured distribution of local polarities and velocities.

### *In silico* experiment: Random polarity field

The goal of this section is to suggest an experimental setup that should permit the antagonism between the different forces in the system due to drag and motor-mediated interactions to be explored. To this end, we performed an *in silico* experiment intended to closely emulate the situation in experiments with *in vitro* filament gels. Photo-bleaching experiments have proven to be a feasible option to simultaneously determine sliding velocities and local gel polarity in filament gels ^17, 18, 31^. In these experiments, the fluorescently labelled microtubules in the gel are photo-bleached along a line by laser light. Due to the motion of the filaments in the gel, the bleached line splits into two lines that move to the right (left) and correspond to left-oriented (right-oriented) microtubules, respectively. From the motion of the two lines, the local velocity and the local polarity can be inferred simultaneously: The local velocity of the left-oriented (right-oriented) microtubules is directly obtained from the velocity of the respective line. Furthermore, the local polarity is determined from the ratio of the bleach intensities of the two lines. The data so obtained only contains local information about the velocity and polarity but no spatially resolved information. In order to make experimentally testable predictions, our goal is, therefore, to derive a relationship between the distribution of measured local polarities and the distribution of measured velocities for which spatial resolution is not necessary.

### Setup of the *in silico* experiment

To illustrate how a given polarity distribution affects the velocity distribution, we consider a specific example, namely a polarity “environment” resulting from random filament assemblies; for details please refer to Appendix “*In silico* experiment: Random polarity field”. We assume that the filament network is nematically ordered and filaments are randomly oriented to the left or to the right, and therefore neglect the possibility that in the experimental system the spontaneous self-organization into the nematic state might involve some polarity sorting. More specifically, filaments are randomly placed in a chamber of size *S* ≫ L with periodic boundary conditions. Since for random filament assemblies there is no reason why the average number of left- and right-pointing filaments should differ, we choose the number density for both left- and right-pointing microtubules to be identical: *μ*^(+)^ = *μ*^(−)^ = *μ*. Importantly, due to the finite extension of the microtubules, the polarity at different positions is not independent. Instead, one finds a positive covariance for the polarities at distances less than one microtubule length *L* apart (Appendix “*In silico* experiment: Random polarity field”). As a result, the polarity profile is not completely random but correlated on lengths smaller than the microtubule length *L* (for a typical profile please refer to Appendix “*In silico* experiment: Random polarity field”).

### Signature of the ambient polarity in the velocity distribution

Based on our theoretical understanding, we expect that, depending on the characteristic length *l*_*c*_, the distribution of velocities is squeezed compared to the polarity distribution. This is because, depending on the ratio of the antagonistic forces, filament motion arises from averaging the polarity over longer (large *l*_*c*_) or shorter (small *l*_*c*_) distances. As we expect the degree of averaging to be reflected in the distribution of velocities, the standard deviation of the microtubule velocities should be an interesting quantity to look at in experiments.

In order to predict the variance of the velocities (ambient polarities) analytically, we describe the local polarity field resulting from the random placement and orientation of filaments in the “experimental” chamber by a set of correlated random variables (see Appendix “*In silico* experiment: Random polarity field”). Using their correlation structure, we average the local polarity according to the expression for the ambient polarity (Eq. 5b) and find (see Appendix “*In silico* experiment: Random polarity field”)

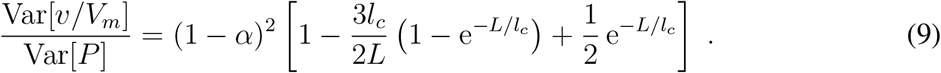

Here, Var[*P*] = Var[*P*(*x*)] = 〈*P*(*x*)^2^〉 − 〈*P*(*x*))^2^〉 denotes the variance of the local polarity, and Var[*v/V*_*m*_] = Var[*v*(*x*)*/V*_*m*_] the variance of the (normalized) velocity *v/V*_*m*_ measured in units of the motor velocity. The above equation implies that the variance of the normalized velocity can be considerably smaller than the variance of the spatial polarity profile; see Fig. 5B. The ratio between the two only depends on the characteristic length *l*_*c*_/*L* and quickly decays with respect to it. For larger *l*_*c*_/*L*, the ambient polarity corresponds to an average over a larger region in space. Therefore, its variance decreases. Due to the linear relationship between the velocity and the ambient polarity, the variance of the velocity decreases to an equal extent.

**Figure 5:**
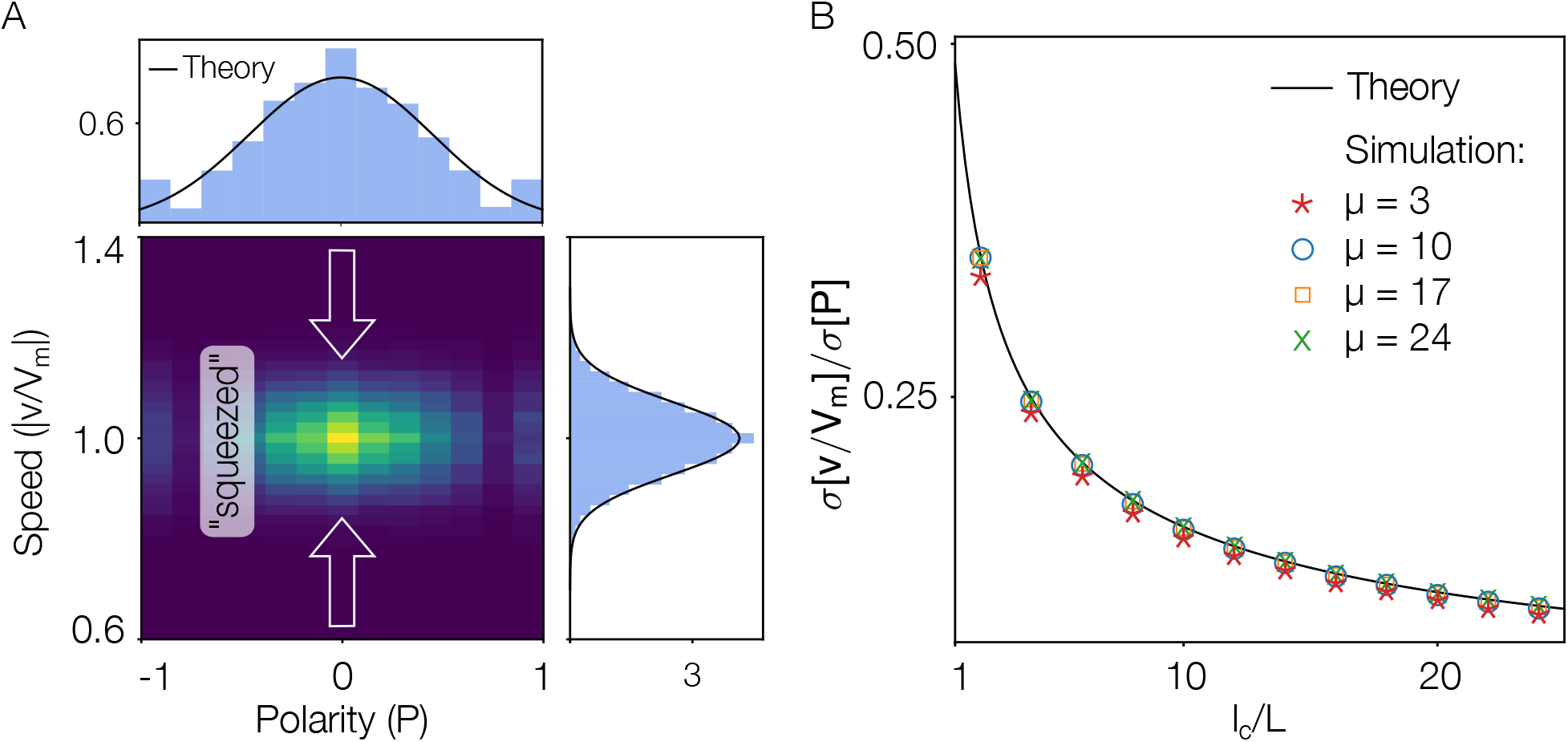
Results for the *in silico* experiment. (**A**) Density plot displaying the probability distribution for all combinations of local polarity, *P*(*x*), and speed, *|v*(*x*)*/V_m_|*, as measured in the *in silico* experiment described in section “*In silico* experiment: Random polarity field”. The histograms at the top and on the right are projections of the density plot on the respective axis. In both cases, the solid line is the corresponding analytic prediction that was obtained by approximating the distributions by a normal distribution with the respective predicted mean and variance. In comparison to the local polarity (top), the velocity distribution (right) is less broad, i.e., it exhibits a smaller but non-zero standard deviation. The parameters are chosen to match the stochastic agent-based simulation described in section “The agent-based model can describe the weak velocity-polarity sensitivity”, namely *μ* = 3 and *l*_*c*_/*L* = 10. (**B**) Ratio between the standard deviations of the normalized velocity, *σ*[*v/V*_*m*_], and of the local polarity, *σ*[*P*], plotted against normalized characteristic length, *l*_*c*_/*L*. The results of the *in silico* experiments (symbols) for *μ* = {3, 10, 17, 24} (red stars, blue circles, yellow squares, green crosses) collapse onto one master curve. The solid line corresponds to the analytic prediction of the master curve. For larger characteristic length, the standard deviation of the velocity decreases relative to the standard deviation of the local polarity. Note that for small *μ* = 3, there is a slight deviation from the master curve. In this case, the variance of the local polarity is so high that the corresponding, approximate normal distribution has not decayed to zero at *P* = ±1 (see histogram at top of A).

In order to compare our results to *in vitro* experiments we assessed the values for both the one-dimensional number density of filaments 2*μ* and the characteristic length *l*_*c*_. From recent experimental data ^18^, we estimated 2*μ* = 6 and *l*_*c*_/*L* ≈ 10; see also section “The agent-based model can describe the weak velocity-polarity sensitivity”. Given these estimates, our theory yields a standard deviation of the polarity distribution 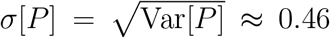, corresponding to a broad range of observable polarities similar to what is seen in experiments. Using our theoretical results we predict the ratio between the standard deviations of the local polarity and the normalized velocity to be approximately 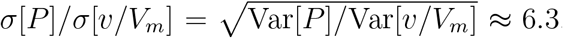. Thus, we expect the mismatch between the widths of the two distributions to be clearly visible in experiments.

### Polarity and velocity distribution in the *in silico* experiment

Figure 5A shows a comparison of the distribution of the local polarity and velocity, as measured in the *in silico* experiment (density plot and histograms) and as predicted analytically (black lines). The density plot shows the measured probability distribution for all combinations of local polarity and velocity. The histograms for both quantities were obtained as projections of the density plot onto the respective axis. While the local polarity takes values in a broad range between ±1 (histogram at top of Fig. 5A), the distribution of the velocity is squeezed to values of approximately (1 ± 0.2)*V*_*m*_ (histogram at right of Fig. 5A).

The disparity between the two distributions nicely illustrates the filtering of high-frequency modes discussed in the section “Analytic solution for motor-induced filament movement”. This filtering is due to long-range interactions induced by the averaging of the polarity field over a length *l*_*c*_. Since the filtering strongly depends on the characteristic length *l*_*c*_, the ratio between the standard deviations of the local polarity and velocity distributions, *σ*[*P*]*/σ*[*v/V*_*m*_], decreases as *l*_*c*_ increases (Fig. 5B). It would be interesting to test this prediction experimentally by changing, for instance, the concentration of the molecular motors, or the drag in the fluid.

Figure 5B shows how the standard deviation of the velocity distribution, normalized to the standard deviation of the local polarity distribution, depends on the characteristic length *l*_*c*_. For small *l*_*c*_ ~ *L*, the effective interaction range of microtubules *l*_*c*_ is small and the microtubule dynamics is predominantly determined by the local polarity at their respective position. Conversely, in the limit of large *l*_*c*_, the dynamics of all microtubules is determined by the same average global polarity. Consequently, all microtubules then exhibit the same velocity and the standard deviation of the velocity *σ*[*v/V*_*m*_] decays to zero. Notably, the normalized curves for different values of the microtubule density, *μ*, collapse onto one master curve when plotted against *l*_*c*_/*L* (see Appendix “*In silico* experiment: Random polarity field”). Thus, in our thought experiment, where we make a certain assumption with regard to the spatial polarity profile, knowledge of the microtubule density is not necessary.

### Experimental relevance

In an experimental filament gel other factors also influence filament dynamics. For instance, as molecular motors randomly attach and detach from microtubules, even microtubules at the same position can interact with a different set of microtubules and thus experience different environments. As a result, different microtubules at the same position might actually have a (slightly) different velocity. Correspondingly, for two experimental realizations with an identical polarity profile, the respective average filament speeds at one position *x* might indeed differ. This effect is not captured by our continuum description, which assumes deterministic velocity profiles *v* ^(±)^(*x*). Thus, we expect a broader distribution of velocities for *in vitro* measurements compared to our theoretical prediction. To gauge the strength of this effect, we compared our theoretical predictions with the results from stochastic agent-based simulations of the system (for details see Appendix “Stochastic, agent-based simulation of the *in silico* experiment”). We find that the specific value of the width of the velocity distribution depends on details of the velocity measurement in the experiments. Nevertheless, irrespective of these details, the velocity distribution is significantly smaller than the width of the polarity distribution.

The *in silico* experiment considered here clearly simulates an idealized system insofar as we have assumed that there is no overall spatial structure. The analysis can be readily extended to a broader class of systems, in which knowledge of the covariance structure of the polarity field (Cov[*P*](*x*, *y*)) is sufficient to predict the covariance structure of the velocity field (Cov[*v/V*_*m*_](*x*, *y*)) (see “Extension of the *in silico* experiment to a broader class of systems”). Since this signature of our results is strongly dependent on the characteristic length, *l*_*c*_, we expect such measurements to provide insight into network parameters. Actually, even low-resolution information on the spatial variation of the polarity field could be helpful to test our predictions. As we have seen above, the Fourier coefficients are suppressed by 1/(1 + (2*πl*_*c*_*k/S*)^2^), *k* ∈ ℤ, in a finite system of size *S* (or, equivalently, by 1/(1 + (*l_c_k*)^2^), *k* ∈ ℝ, in the infinite system). So, for large *l*_*c*_, the velocity modes with wave vector *k* ≥ 1/*l*_*c*_ should not be visible in experiments.

## 3 Summary and discussion

In this work, we have considered a mesoscopic model for microtubule dynamics in a nematic, motor-crosslinked network. So far, research has focused on either the dilute or heavily crosslinked limit. Strikingly, the observed behavior in these two cases is qualitatively different: While in the dilute case the microtubule velocities strongly depend on the local network polarity ^20, 21^, in the heavily crosslinked case the velocity has been found to be independent of the polarity ^14, 16, 18^). These distinct phenomenologies are puzzling, as the underlying microscopic motor-mediated microtubule interactions are presumably the same in both cases. Starting from these filament interactions, we have shown how the interplay between movement resulting from motor-crosslinking and the countervailing effects of fluid drag determines the sensitivity of the local filament dynamics to the network polarity. Thereby we provide a better understanding of the essential physical principles that lead to such diverse dynamics.

To this end, we derived a non-local mean-field theory of our system from the microscopic interactions. This theory enabled us to obtain an explicit analytic expression relating the local microtubule velocity to the spatial polarity profile. Our key result is that the local velocity depends on the local *ambient polarity*, which is given by the averaged polarity a microtubule senses in its environment. More specifically, the local velocity is given by the convolution of the polarity and an exponentially decaying interaction kernel with *characteristic propagation length*, *l*_*c*_. Hence, it is not the local polarity at the position of a microtubule that determines its motion but rather the entire polarity profile in an environment of length *l*_*c*_. This finding implies that a one-to-one mapping between the local velocities of microtubules and the local polarity as shown in Fig. 2 is not the whole story. Instead, in order to predict the velocity at a specific location, knowledge of the spatially varying polarity profile in the entire vicinity is needed. In general, such detailed spatial information appears to be inaccessible with current experimental techniques. Fortunately, in order to infer the distribution of velocities from the distribution of local polarities, such detailed information is not essential. For example, in a gel where microtubules are randomly placed in an experimental chamber and stochastically oriented, our theory predicts how the variances of the local polarity and of the velocity are related.

The relationship between the velocity and polarity distributions strongly depends on the characteristic propagation length *l*_*c*_, which is an important emerging length scale in the system. It can be interpreted as a non-local interaction range of filaments, and is determined by the ratio between the average motor-driven force on a microtubule and the microtubule’s drag in the fluid. Thus, this intrinsic length reflects how effectively motion generated at one position is propagated through the interconnected network of filaments. It strongly depends on the network properties.

We have identified a common mechanism explaining the microscopic origin of both uniform filament motion in percolated nematic networks and the strong polarity dependence of microtubule motion in dilute systems: Due to their finite extension, microtubules directly interact with several parallel and antiparallel neighbors within a spatial ranges equal to twice their filament length. Motors between parallel microtubules induce a resistance against relative motion and thus promote uniform motion of crosslinked microtubules. Thereby, motion generated by antiparallel interactions translates through the percolated network of microtubules even into regions with only parallel and no antiparallel interactions where *a priori* no motion is expected. The degree of efficiency of this propagation of motion is quantified by the characteristic propagation length *l*_*c*_. Hence, it is influenced by the average number of motors per interaction and the drag of filaments in the fluid, among other factors. Filaments at distances larger than *l*_*c*_ apart can be considered to be part of disconnected patches. That is, for small *l*_*c*_ only motor-crosslinks between nearest neighbor filaments are relevant for filament motion, as in the dilute limit. For this case, we recover the linear relationship between local polarity and filament velocity ^19–21^. On the other hand, in the limit of large *l*_*c*_, which corresponds to systems where the patch size exceeds the system size, we find a dependence of the velocity on the global polarity only. Here, the velocity for equally oriented microtubules is the same everywhere in space. In particular, our results explain the weak sensitivity of the filament velocities to the local polarity observed in recent experiments ^18^ and in the spindle apparatus ^11, 14, 16^.

In particular we predict a strong dependency of the velocity distribution on the characteristic propagation length. In order to test this prediction, we suggest a practicable *in vitro* experiment whose feasibility we assessed with the help of an *in silico* experiment intended to mimic the suggested *in vitro* experiment. Intriguingly, it is not necessary to determine the entire spatial polarity and velocity profile to check the validity of our theory. Instead, it suffices to determine the polarity and velocity distributions by measuring the local velocity and polarity at random positions in the filament gel. When plotting the ratio of the standard deviations of the polarity and velocity distribution against the characteristic length *l*_*c*_, we expect the data to collapse onto a master curve, irrespective of the explicit number of filaments in the experimental chamber (Fig. 5). Furthermore, the ratio of the standard deviations of the polarity and velocity distributions for a specific experimental setup could be used to identify the characteristic propagation length *l*_*c*_ and, allow one to draw conclusions regarding network features (Eq. 6).

Microtubule motion in mitotic spindles formed in Xenopus egg extract is a prominent example for polarity-independent sliding. The polarity profile in these spindles is approximately linear, ranging from zero polarity in the center to highly polar regions at the spindle poles ^17^. Nonetheless, microtubules drift with roughly constant velocity towards the spindle poles, especially if dynein is inhibited. Our theory can account for this behavior. In particular, the individual velocities deviate only slightly from the mean velocity if motor-crosslinking is strong, i.e. if the characteristic length exceeds the system size (see Sec. “A simple example: The linear polarity profile”). Interestingly, for biologically plausible parameters the interaction range is of the same order as the length of the spindles formed in Xenopus egg extracts, *l*_*c*_ ∝ 30 − 80 μm. Correspondingly, as seen in Fig. 4(A), the velocity of the poleward moving microtubules is expected to be slightly smaller close to the pole than in the center of the spindle. This variation is due to the dependence of the velocity on the ambient polarity (the local polarity environment). Taken together, our results suggest that, depending on the value of the characteristic length compared to the spindle size, the spatial polarity profile and, in particular, the fact that the poles are highly polar, could be significant for the velocity profile as well. To examine this behavior experimentally, it would be instructive to investigate the velocity distribution of microtubules in a dynein-depleted, unfocused spindle as a function of the distance from the spindle boundary.

From a broader perspective, it would be interesting to extend our work on nematic networks to a more general description of filament gels. To this end, it could be promising to start from recent work on heavily crosslinked filament gels, where a sophisticated hydrodynamic framework has been established from microscopic properties ^18^. This theoretical framework assumes an infinitely large characteristic length *l*_*c*_, so that motion generated at one position propagates through the whole network without loss. Our results suggest that incorporating an exponential interaction kernel into this framework can provide a more comprehensive description of filament motion in crosslinked gels. Such a description would also offer the chance to understand the transition from heavily crosslinked to weakly coupled gels.

## 4 Methods and Materials

### Numerical simulation Agent-based simulation

To simulate the filament gel, we implement an agent-based simulation consisting of *M*_*l*_ left- and *M*_*r*_ right-oriented filaments. As described in detail in section “*In silico* experiment: Random polarity field”, we randomly place the filaments in a one-dimensional box with periodic boundary conditions. Next, a vector 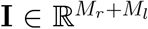 consisting of the numbers of overlapping filaments for each filament *i* is generated. From this vector, a probability vector 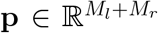 is derived so that the average number of interaction partners per filament is given by *N* = *I*_*i*_ *p*_*i*_. Out of the *I*_*i*_ possible interaction partners of filament *i*, we accept an interaction with probability *p*_*i*_ and reject it with probability 1 − *p*_*i*_. Once the interactions are determined, we construct a set of *M*_*l*_ + *M*_*r*_ coupled linear equations on the basis of the force balance equation

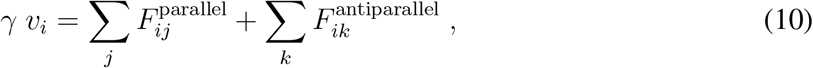

and weigh each interaction by the overlap between the filaments. Here *j* runs over the parallel interaction partners and *k* over the antiparallel interaction partners of filament *i*. The velocities of each filament *i* are then obtained using matrix inversion.

### Continuum simulation

For the continuum simulation, we generate a polarity profile analogous to that in the agent-based simulation. Then we use our theoretical results (Eqs. 5a, 5b) but perform the integration numerically to obtain the velocity field.

### Analytical description

Our analytical description starts from the microscopic linear force-velocity relations describing the force exerted by two individual parallel or anti-parallel microtubules on each other, Eqs. 1. Then, instead of looking at all individual microtubules and interactions, we move to another level of description. Microtubules and their velocities are described in terms of spatially dependent density and velocity fields. Furthermore, the molecular motors mediating the forces between the microtubules are only taken into account implicitly, in accordance with a uniform spatial density. As a result, the interaction strength between two overlapping microtubules is proportional to the overlap region. Taken together, these assumptions yield Eq. 3. Solving it for the velocities leads to the main result, the velocity-polarity relations given in Eqs. 5a and 5b together with the definition of the characteristic length, Eq. 6. For details and the two applications of our main result, we refer the interested reader to the Appendix.

## 5 Acknowledgments

We thank Silke Bergeler, Philipp Geiger, Emanuel Reithmann and Patrick Wilke for helpful feedback on the manuscript. This research was supported by the German Excellence Initiative via the program “NanoSystems Initiative Munich” (NIM). I.R.G. is supported by a DFG fellowship through the Graduate School of Quantitative Biosciences Munich (QBM). We also gratefully acknowledge financial support by the Deutsche Forschungsgemeinschaft (DFG) via Collaborative Research Center (SFB) 863.

## Appendix

## S1 A continuum model for motor driven filament motion

Here we derive a solution for our continuum model of filament motion. As a starting point, we use the coupled set of integral equations, Eqs. 3, in the main text which read

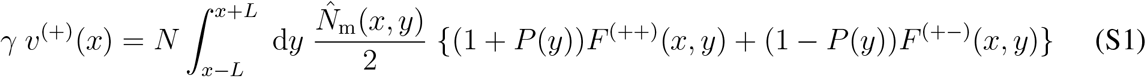

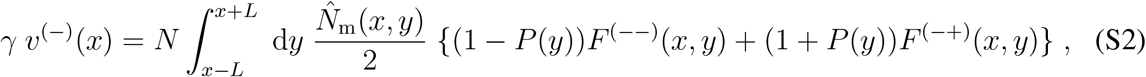

where we used *φ*^(±)^ = 1/2(1 *± P*(*y*)).

In general, it is quite challenging to provide an analytic solution to a set of coupled integral equations. Here, however, one can make use of the fact that the difference of the velocities, *v*^(+)^ − *v*^(−)^, takes a quite simple form. Namely,

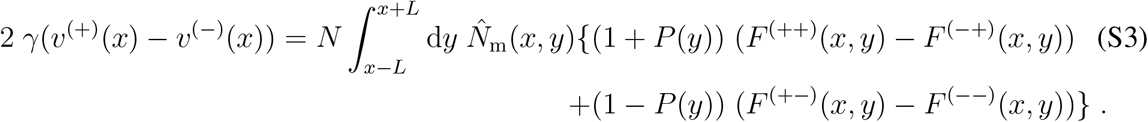

The difference of the contributing forces reads

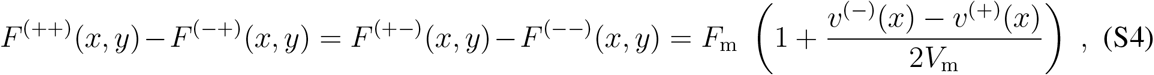

and is a function of *x* only.

Substituting Eq. S4 into Eq. S3 and performing the integration yields

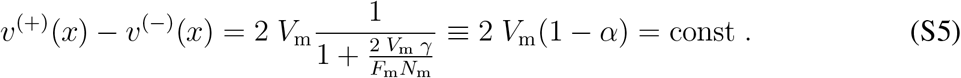

As a result, *v*^(−)^ is expressed in terms of *v*^(+)^, and we can use this relation to decouple Eq. S1 and Eq. S2.

The resulting integral equation reads

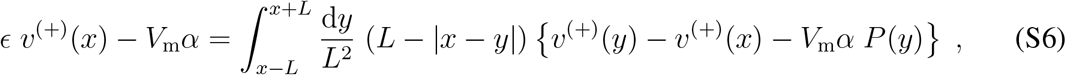

where we introduced 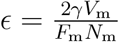.

To proceed further, we rewrite *v*(*y*) and *P*(*y*) in terms of their full Taylor expansions around *x* and shift *v*^(+)^ by *α/ϵ V*_m_, i.e., *v*^(+)^ → *v*^(+)^ − *α/ϵ V*_m_.

Performing the integration yields an ODE coupling the velocity to the polarity field. It reads:

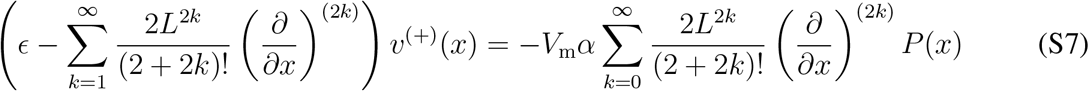

## S2 An analytic solution for filament motion in a nematic network

To find a feasible expression relating the velocity and polarity field, we apply the Fourier transformation to Eq. S7. Our system – recast in *v*^(+)^(*k*) and *P*(*k*) – becomes

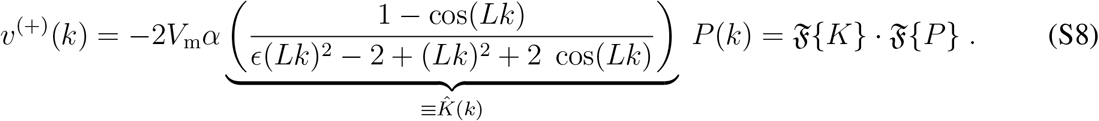

where 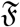 denotes the Fourier transformation operator. From the convolution theorem, we directly find that *v*^(+)^(*x*) is given by the convolution of *K*(*x*) and *P*(*x*). So, in order to tackle our original equations, we are left with finding the Fourier back transformation of 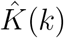, i.e., we need to solve

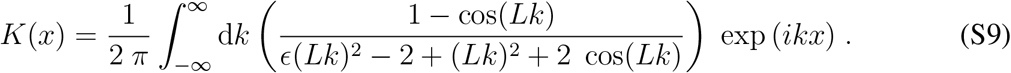

To proceed further, we assume that the integral can be performed using the residue theorem, i.e.,

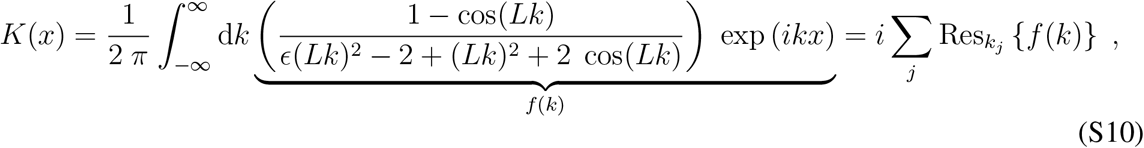

with the sum running over all poles in the upper half plane (lower half plane) if *x* > 0 (*x* < 0). In the following, we will restrict the discussion to the case *x* > 0 since the calculations for *x* < 0 are analogous. Note that we exclude the case *x* = 0 explicitly, and assume a smooth solution at *x* = 0. For simplicity, we will use dimensionless variables in the following and recast *k* → *kL* and *x* → *x/L*.

To find the potential residues, we search for all solutions of the equation

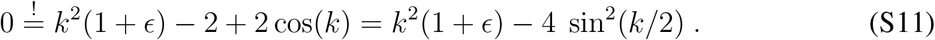

in the complex plane. In the following, we will use *k* = *a* + *ib* with *a*, *b* ∈ ℝ. Using this notation, the problem of finding possible residues of *f*(*k*) has shifted to finding solutions to the equations

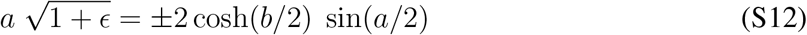

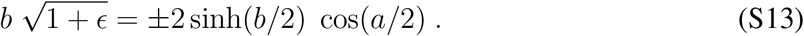

We split the discussion into (*i*) real, (*ii*) imaginary and (*iii*) complex solutions of Eq.S11. *a*^*^, *b*^*^ will denote solutions of the above equation system.

i. **real solutions** (*b*^*^ = 0) The only real solution of Eq. S11 is given by *a*^*^ = 0.
ii. **imaginary solutions** (*a*^*^ = 0) Eq. S12 is always fulfilled for 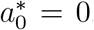. Moreover, Eq. S13 always has two solutions for *ϵ* > 0 which can be estimated as 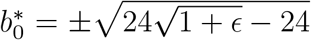 for small *ϵ*.
iii. **complex solutions**(*a*^*^ ≠ 0, *b*^*^ ≠ 0) The complex solutions are not trivial to find. First, note that the discussion can be split into positive and negative signs of the right-hand side of Eq. S12 and S13.

For a negative sign on the right-hand side, if *a, b* > 0, cos(*a*/2) and sin(*a*/2) have to be negative for the equation system to be solvable, i.e., we know *a* ∈(2*nπ*, (2*n* + 1)*π*) with *n* = 1, 3, 5….

Investigating Eq.S13 alone, the minimal *b*^*^(*a*) is found for *a* = 2*πn* and can be estimated as 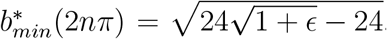, i.e.,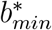 is close to 0 for small *ϵ*. Moreover, one finds that *b*^*^(*a*) is monotonically increasing in the interval *a* ∈ (2*nπ*, (2*n* + 1)*π*) and *b*^*^(*a* → (2*n* + 1)*π*) → *∞*. Equation S12 implies that cosh(*b/*2) has to be sufficiently large if *a*∈(2*nπ*, (2*n* + 1)*π*) with *n* = 1, 3, 5…. Especially for large *n* this means that *a*^*^ has to be close to (2*n* + 1)*π* with *n* = 1, 3, 5…, (i.e., 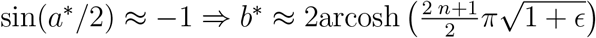. Taken together, this yields

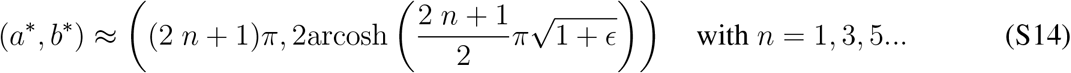

A similar argumentation for a positive sign on the right-hand side of Eq. S12 and Eq. S13 and *a, b* > 0 yields the same result for *n* = 2, 4, 5…. The cases *a* > 0, *b* < 0, *a* < 0, *b* > 0 and *a* < 0, *b* < 0 are analogous.

Taken together, we find the solutions

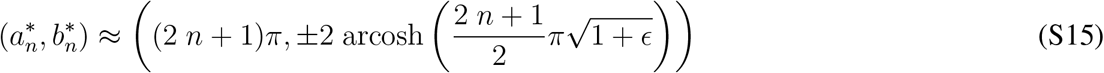

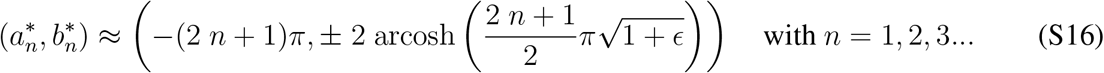

Figure S2 shows a comparison between the numerically found roots and our approximation for *ϵ* = 0.1.

Since there is an infinite number of poles with arbitrarily large real part, one can not proceed as usual and use the residue theorem without any additional considerations. So, to make further progress, we continue as follows: First, we show

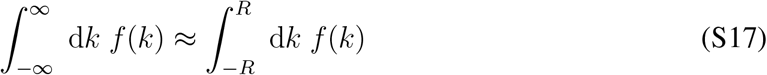

for sufficiently large *R*.

Second, we compute the integration along the path 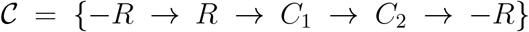 according to the residue theorem:

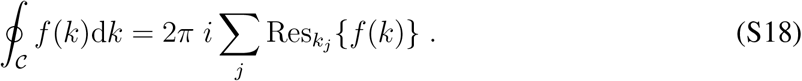

**Figure S1:**
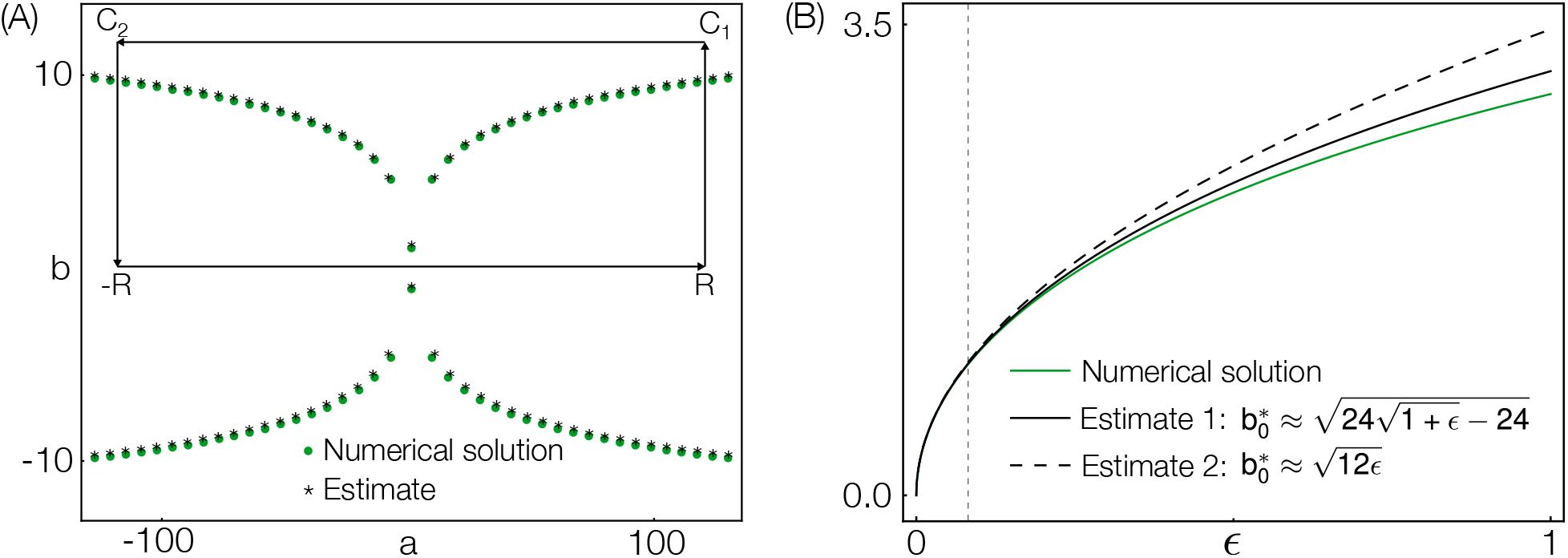
(**A**) Comparison of the estimate of the solutions of the equation system Eq.S12,S13 with the solutions obtained numerically (the results are compared exemplary for *ϵ* = 0.1). (**B**) Comparison of two estimates for the imaginary pole with the corresponding numerical solutions. The gray dashed line indicates the *ϵ*–value for which *l*_*c*_ ≈ *L*.

Third, we show that the path in the complex plane gives a vanishing contribution. Then the real Integra 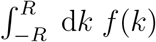 can be estimated by

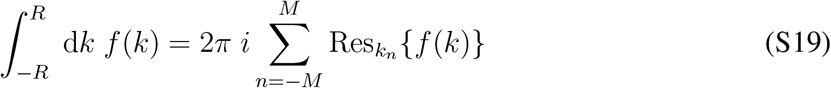

where *k*_*n*_ = *k*_−*M*_ …*k*_*M*_ denotes the poles in the interior of the integration path sorted by increasing real part. The notation is chosen so that *k*_0_ denotes the purely imaginary pole.

The first step is straightforward since

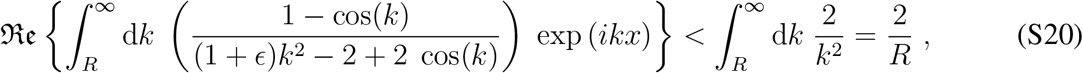

i.e.,

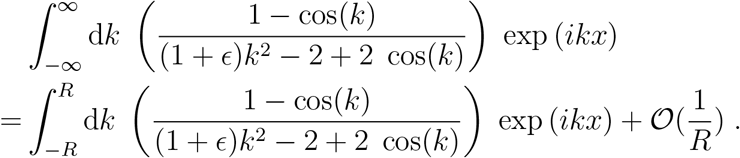

Next, we aim on showing that there is a path *R* → *C*_1_ → *C*_2_ → −*R* so that the integration along that path gives zero contribution in the limit of large *R*.

Usually, if one deals with functions of the form *g*(*k*) exp(*ixk*) one can make use of Jordan’s lemma which states that the integration along the semicircular contour in the upper half plan (lower half plan) for *x* > 0 (*x* < 0) aims to zero for the radius *R* ≡ |*k*| → ∞. However, if *f*(*k*) is rewritten in the form of *g*(*k*) exp(*ixk*) with *g*(*k*) = (1 − cos(*k*))/((1 + *E*)*k*^2^ − 2(1 − cos(*k*))), we face the problem that *g*(*k*) does not converge uniformly to zero since *g*(*k*) has an infinite number of poles in the upper (and lower) half plane. So, to proceed further we aim on proving that there exists a path so that the contribution of the contour in the complex plane goes to zero for large radius.

First, note that

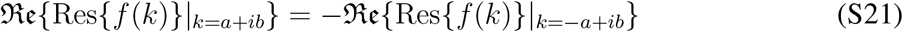

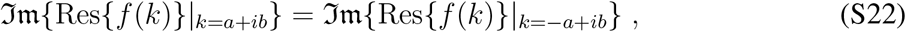

i.e., if the integration path is chosen in a way to symmetrically include the poles in the upper left and upper right quarter we know 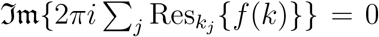. Hence, if the contour integral (Eq. S18) is calculated explicitly along such a path, the result has to be real. In the following, we will make use of this fact. To prove that the integration along the contour in the complex plane goes to zero (for large *R*), we choose an explicit path along a rectangle as shown in Fig.S2. More concretely, the vertices are defined by *R* = 2*nπ*, *C*_1_ = 2*nπ* + *i*2*nπ* and *C*_2_ = −2*nπ* + *i*2*nπ* for *n* ∈ ℕ. Since the overall contour integral is real, we only care about terms which can give a real contribution, i.e., we investigate the real (imaginary) parts of the integrand for the integration paths parallel to the real (imaginary) axis:

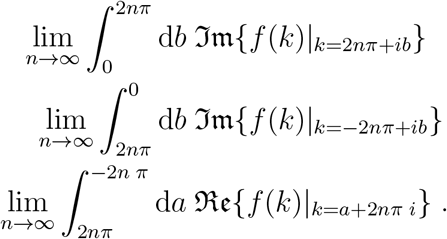

To prove that those terms give zero contribution for *n* → ∞ (and thereby *R* → ∞), we aim on finding an integrable majorant of the above expressions, and then swap the integration and the limit. It is possible to show that

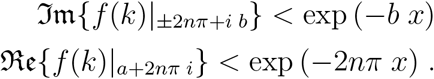

Thus, we found an integrable majorant for both paths. Furthermore, since 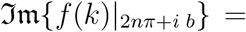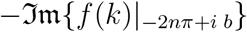, the discussion can be restricted to one of the paths *R* → *C*_1_ or *C*_2_ → *−R.* Moreover, as we found an integrable majorant for the parts of the contour in the complex plane, we can swap the integral and the limit. Since 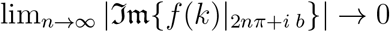, the paths *R* → *C*_1_ and *C*_2_ → *−R* give zero contribution to the contour integral. The same holds true for the integration along the path *C*_1_ → *C*_2_. Thus, the integration along the contour in the complex plane gives zero contribution for *n* → ∞ and we can calculate the desired integral by

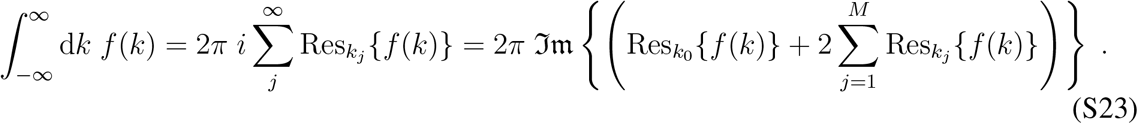

In the last step, we made use of Eq.S21. In the following, we will denote the contribution of the *k*_0_ residue to the integral as *f*_0_(*x*) and the contribution of the sum over all other poles as *f*_∞_(*x*) Using the estimated expression for the poles of *f* (*k*) and making use of the fact that 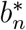 is large, we find an estimate for the imaginary part of the residue which reads

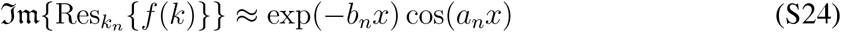

for *n* = 1…*M*. Here, we assumed that *b*_*n*_*x* is sufficient large, i.e., we expect deviations for *x* → 0.

For *k*_0_ we find the residue (for small *ϵ*)

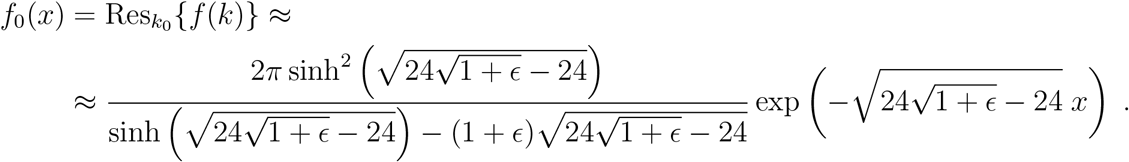

So, to find a closed form of the integral, we are left with finding an expression for the sum

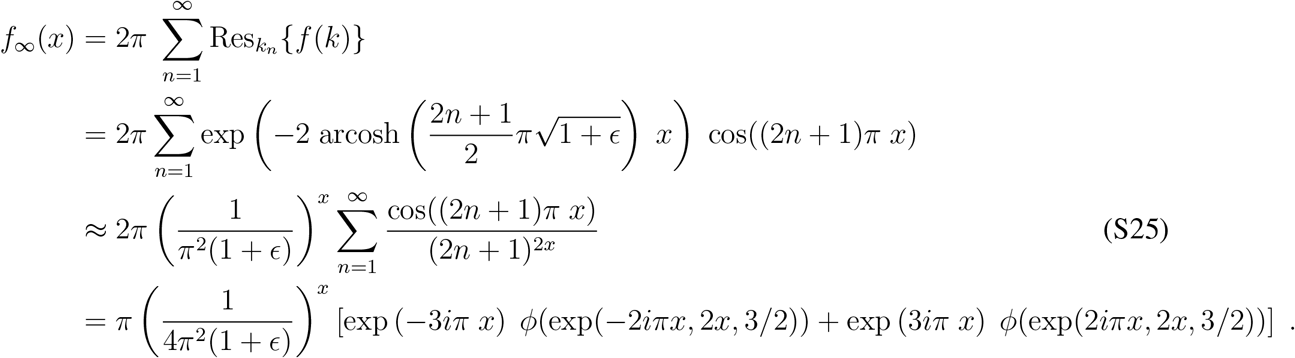

For the approximation we used arcosh(*x*) ≈ ln(2*x*) for large *x*. Here, 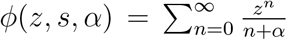 denotes the Lerch zeta function. Fig. S2 (A) shows a comparison between our analytic result and the sum over the first 5000 numerically obtained residues. For a better comparison, we also perform the sum over the approximated residue (Eq.S25) for the first 5000 terms. Moreover, we compute the ratio of *f*_∞_/*f*_0_ to get insight into how much *f*_∞_ contributes to the overall integral. Taken together, this yields the desired expression for the integral and, thereby, an expression for the interaction kernel in real space which reads

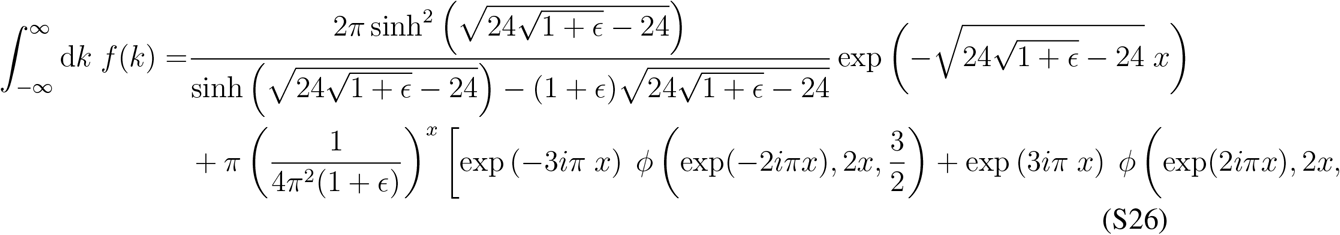

This expression, however, is not particularly intuitive. Therefore, we seek for a simpler and more meaningful expression.

Fig. S2 (B) indicates that *f*_∞_ is not particularly relevant for the overall integral, i.e., it is promising to use only the 0*th* residue to approximate the integral. Moreover, *ϵ* is known to be small for biologically meaningful parameters. In hindsight, one can argue that a value of *ϵ* > 1/12 is not particularly meaningful since for *ϵ* < 1/12 our result suggests an interaction range *l*_*c*_ < *L*, implying that the microtubules interact only over a distance smaller than one microtubule length. Therefore, it makes sense to consider the limit of small *ϵ*. Using only the 0*th* residue and considering the lowest order of *ϵ* yields

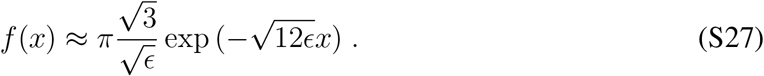

Fig. S2 (B) shows the estimate of the imaginary pole in comparison to the more accurate estimate and the numeric result. Fig. **??** shows a comparison between the numerical solution of the integral, our analytic expression (Eq. S26) and the 0*th* residue approximation for small *ϵ* (Eq. S27).

Finally, making use of the convolution theorem, Eq. S27 and Eq. S8 yield the desired expression for the filament velocity. Going back to natural variables, i.e., *x* → *x L* and 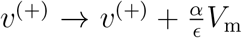 gives the expression used in the main text:

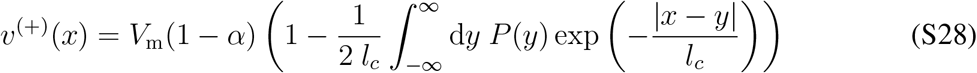

where we introduced 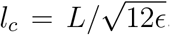. The occurrence of the absolute value is due to the fact that an integration over the lower half plane gives an analogous result as compared to the one for the upper half plane.

## S3 Fourier representation of the ambient polarity

The ambient polarity is given by the convolution of the local polarity with an exponentially decaying interaction kernel, Eq. 5b. As the Fourier transformation of a convolution of two functions is given by the product of the two Fourier transformations, it is instructive to consider the Fourier representation of the polarities. We distinguish two cases. First, we consider an infinite system where the fields are defined on the entire real axis, and, second, a periodic system with period *R*.

**Figure S2:**
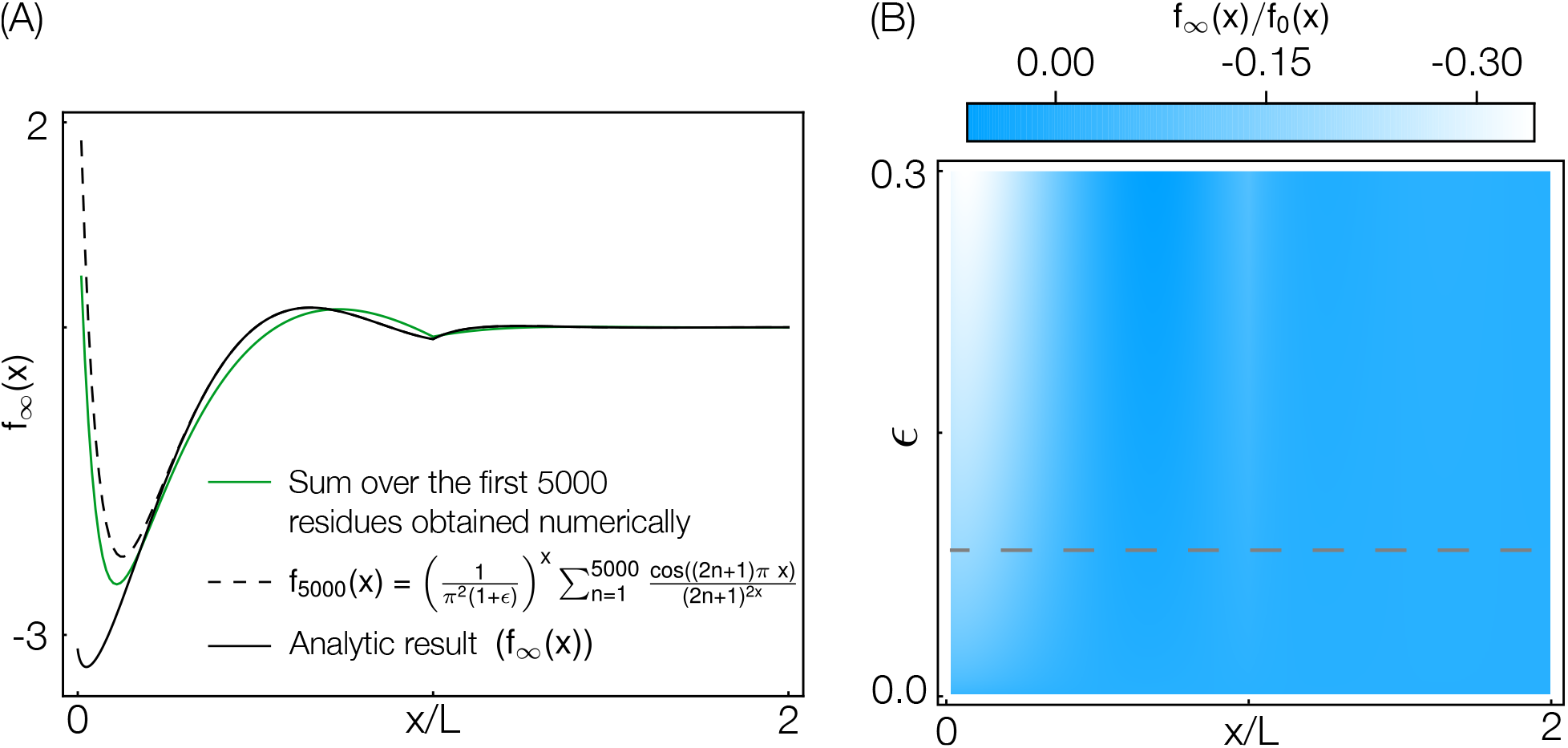
(**A**) Analytic result for the sum over the residues in the complex plane in comparison to the sum over the first 5000 numerically obtained residues. To provide a more accurate comparison, we also compute Eq.S25 for the first 5000 terms. (**B**) To compare the contribution of the *k*_0_ residue and the sum over the other residues to the overall integral, we plot the ratio *f*_∞_(*x*)/*f*_0_(*x*). The dashed line indicates the *ϵ* value where *l*_*c*_ ≈ *L*

**Figure S3:**
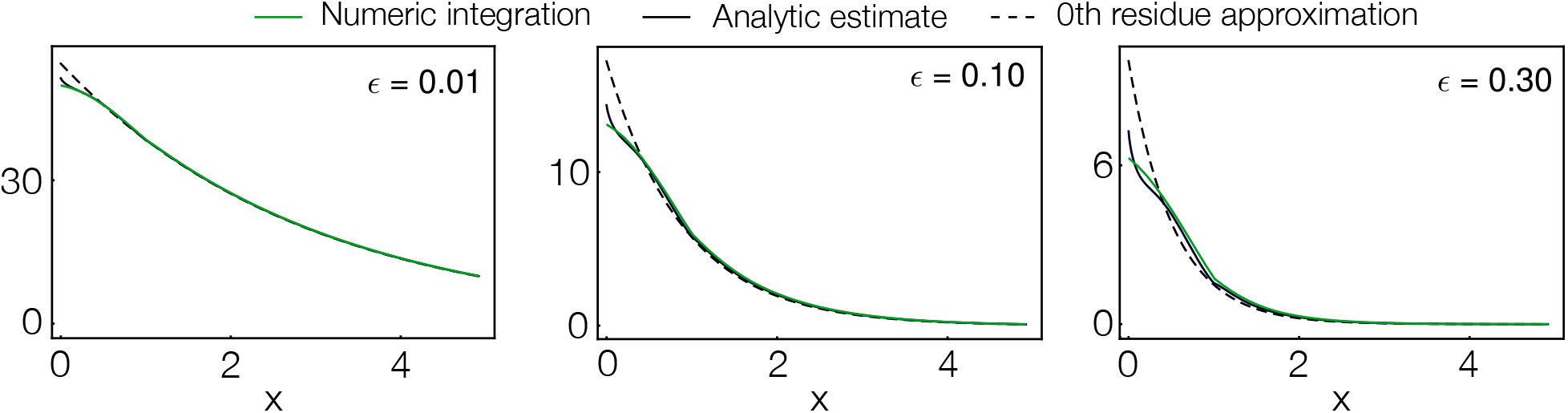
Comparison between numerical integration, analytic approximation and the 0th residue approximation used in the main text. Note that the deviation between the approximations for larger *ϵ* is manly caused by the wrong estimate of the *k*_0_ residue.

## Infinite case

For a polarity, *P*(*x*), defined on the real axis, *x* ∈ ℝ, we define the Fourier transformation as

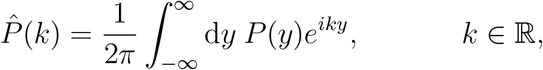

with the corresponding back transformation 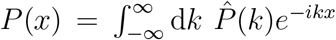. Similarly, the Fourier transformation of the ambient polarity is

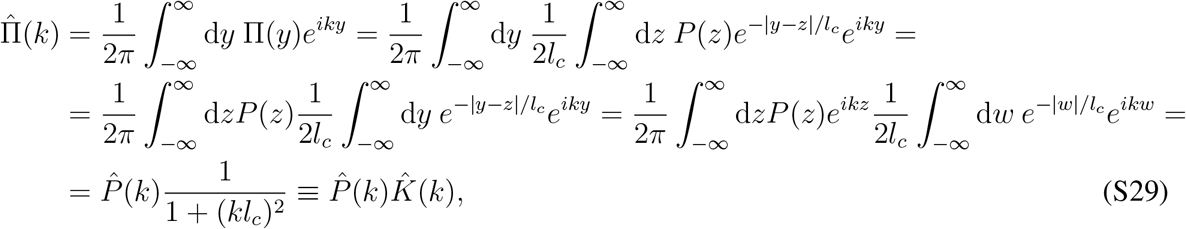

where we exchanged the integrals and performed the Fourier transformation of the exponentially decaying interaction kernel, 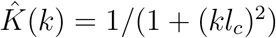. This result implies that the spatial modes are suppressed according to a Lorentzian. So, faster fluctuations are damped more, and the ambient polarity does not exhibit spatial variations corresponding to large wave vectors *k* ≫ 1/*l*_*c*_ (small wavelength). Intuitively, the lack of fast fluctuations in the ambient polarity is due to the averaging of local polarities in a range of size *l*_*c*_. As we will see in the following, we get a very similar result in the periodic case.

## Finite interval with periodic continuation

If the system is periodic with period *R*, the polarity, *P*(*x*), *x* ∈ [0, *R*], is described by a Fourier series, 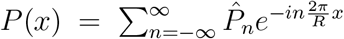, with Fourier coefficients

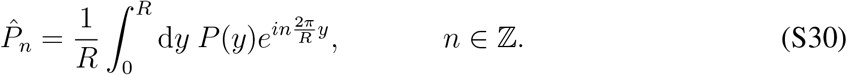

The Fourier coefficients for the ambient polarity, 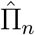, *n* ∈ ℤ are given by

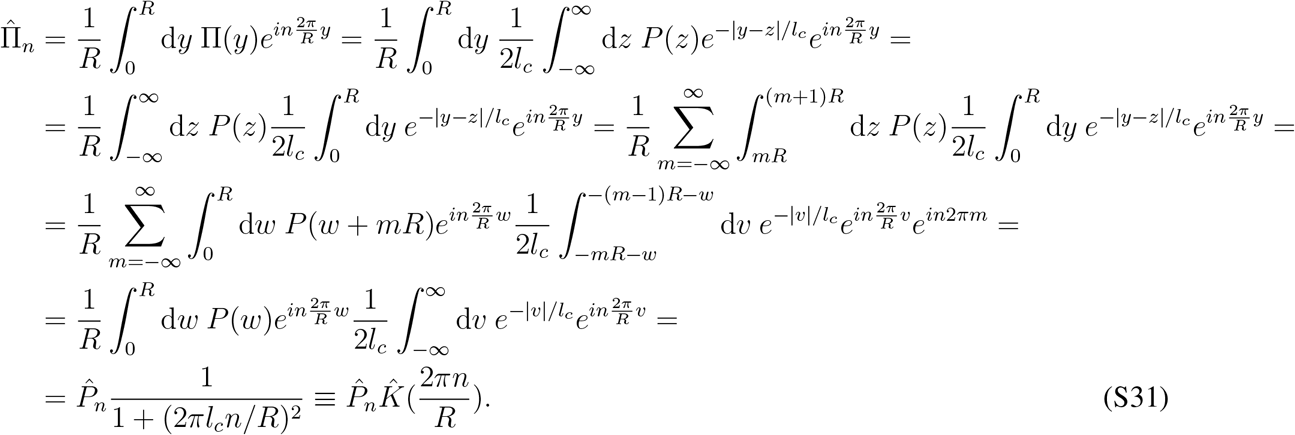

In these steps, we exchanged the integrals and used that the infinite integral can be expressed in terms of an infinite sum of integrals over a period *R* each. Furthermore, we used the substitutions *w* = *z* − *mR* and *v* = *y* − *mR* − *w* and the periodicity: *P*(*w* + *mR*) = *P*(*w*) and *ϵ*^*in*2*πm*^ = 1 for *m* ∈ ℤ. We could have guessed this result from the result of the infinite case, Eq. S29, as in the periodic case only wave vectors *k* which are a multiple of 2*π/R* are possible: *k* = *n*2*π/R* for some *n* ∈ ℤ. So, again, fast fluctuations are strongly suppressed in the ambient polarity.

## Relevance

Importantly, these results are not restricted to a specific class of polarity profiles but generally capture the relationship between the local and ambient polarity in an infinite (large) system. Hence, the ambient polarity (the velocity) is expected to vary at most on length scales larger than the characteristic length *l*_*c*_. Related to this, extreme values of the local polarity are averaged out and do not show up in the distribution of velocities. This observation is illustrated by two examples in the main text, namely the pedagogical case with linear polarity profile and the *in silico* experiment with random polarity profile.

## S4 Linear polarity profile

As a first example to illustrate the relationship between the ranges of local and ambient polarity, we consider a linear polarity profile *P*(*x*) = *a* * (*x − S/*2) on a finite interval *x* ∈ [0, *S*] (where |*a*| ≤ 2/*S* to ensure that the polarity does not exceed ±1). This situation is in contrast to the main analysis in the manuscript which focuses on an infinite system. Consequently, we have to specify some boundary conditions.

## Motivation of boundary conditions

In order to fix the boundary conditions, we start from the premise that even at the boundary, the system is dense and the number of interaction partners of a microtubule is limited by the number of neighbors and not by the overall number of microtubules. In other words, the number of interaction partners per microtubules is the same, irrespective of whether the microtubule is located in the bulk of the system or at the boundary. This implies that microtubules at the left (right) boundary have twice as many crosslinks towards their right (left) as compared to microtubules in the bulk of the system (and none to the left (right) due to the boundary). To approximately account for this effect, we mirror the polarity profile at its boundaries *x* = 0 and *x* = *S*, so we have *P*(*−x*) = *P*(*x*) and *P*(*S − x*) = *P*(*S* + *x*). By repeated application of this mirroring, the polarity profile is continued to the entire real axis. The resulting continued spatial polarity profile is 2*S* periodic. Thus, we approximate our finite system by a periodically continued spatial polarity field with the same (infinite) interaction kernel exp(−|*x* − *y*|/*l*_*c*_) as for the infinite system.

## Fourier coefficients

The local polarity is represented by a Fourier series with Fourier coefficients given in Eq.S30 for *R* = 2*S*. More concretely, due to the symmetry *P*(*x*) = *P*(*−x*) the Fourier coefficients are given by

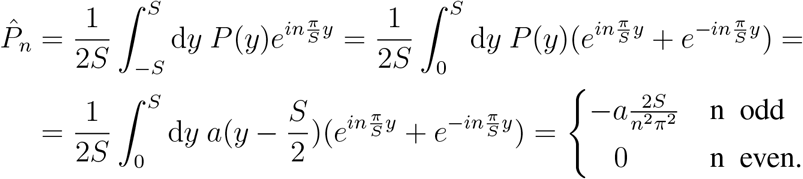

Eq. S31 then implies that the Fourier coefficients of the ambient polarity are given by

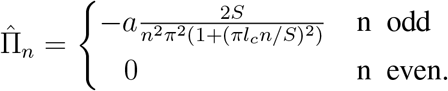

## Ratio between the ambient and local polarity range

From the Fourier coefficients, we can straightforwardly determine the ratio between the ranges of local and ambient polarity. Due to the monotonicity of the local polarity and the from-the-center decreasing interaction kernel, the maximum (minimum) of the ambient polarity is at the same location as the maximum (minimum) of the local polarity. In order to compute the ratio between the two ranges of values, it is thus sufficient to determine the ambient polarity at *x* = *S* where it attains its maximum (due to the symmetry, the minimal value at *x* = 0 corresponds to the inverse of the maximal value):

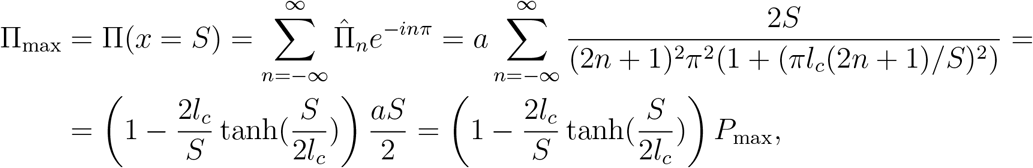

where we used that *P*_max_ = *P*(*x* = *S*) = *aS/*2. A more instructive expression can be obtained by using a large wavelength approximation, describing the local and ambient polarities by their lowest modes *n* = ±1, 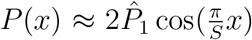 and 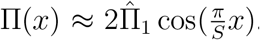. Thereby, the ratio of the ranges of the local and ambient polarity is approximately

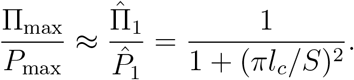

As can be seen in Figure 4, this expression is a fairly good approximation, in particular for large enough *l*_*c*_/*S* as expected. It predicts that the range of the ambient polarity is strongly “squeezed” for large characteristic length.

## S5 *In silico* experiment: Random polarity field

In this appendix, we give details on how we construct the random polarity field in the *in silico* experiment described in section “*In silico* experiment: Random polarity field”. Furthermore, we derive the formula for the ratio between the variances of the local and ambient polarity, Eq. 9.

## Construction

Our goal is to mimic a realistic polarity profile that arises from spontaneous self-organization of stabilized microtubules and crosslinking motors into nematically aligned filament gels in the experiments. To this end, we construct a network of randomly oriented microtubules with approximately constant density. Explicitly, the network is created in the following way: First, we divide the system of size *S*(*S* ≫ *L*) into small containers of size Δ*x*. Second, for each container the numbers of microtubules with midpoint in the container and pointing to the left or right, respectively, is drawn independently according to a binomial distribution with mean *μ*Δ*x/L* for each type of microtubules (left/right). Here, *μ* can be interpreted as the average 1D number density of left- or right-oriented microtubules, respectively.

## Distribution of the local polarity

This procedure results in a spatial profile for the number of microtubule midpoints which is rough. However, due to the finite extension of microtubules the corresponding polarity and density profiles, *P*(*x*) (*ρ*(*x*)), are smoothed. The local polarity at a certain position is determined by all the microtubules that cross this position and not only those whose midpoint is located there: Denote by 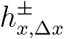 the (randomly drawn) number of left- or right-oriented microtubules with midpoint in [*x, x* + Δ*x*) and by 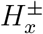 the (resulting) number of left- or right-oriented microtubules that cross position *x*. As all microtubules with midpoints in [*x* − *L/*2, *x* + *L/*2] pass through *x*, these quantities are related as follows:^3^

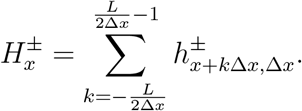

^3^

Here 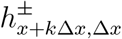 are independent and identically distributed according to a Binomial distribution with mean *μ*Δ*x/L*. For small enough intervals Δ*x*, we can approximate these Binomial distributions by Poisson distributions:

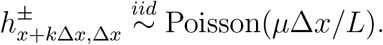

As 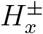 corresponds to the sum over *L/*Δ*x* of these random variables, it is distributed according to a Poisson distribution as well, and the mean is given by the sum of the means:

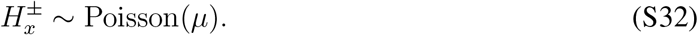

Importantly, the finite extension of the microtubules introduces correlations in the number of microtubules crossing different positions *x*. That is, the quantities 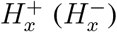 for different *x* are not independent: Their covariance is given by

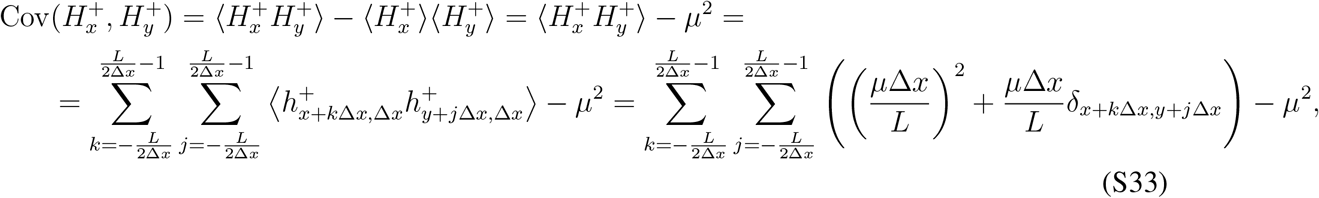

where in the last step we used that the random variables 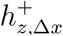 for different *z* are independent with mean *μ*Δ*x/L*: If *z*_1_ ≠ *z*_2_ we have 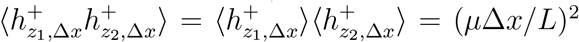. In contrast, the second moment of a Poisson distribution with mean *λ* is given by *λ*^2^ + *λ*, so 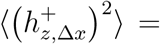 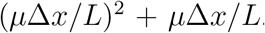. Performing the sum over the constant 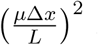 in Eq. S33, we find the following expression for the covariance:

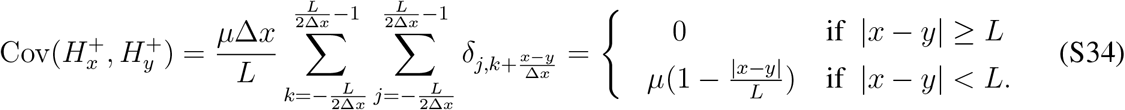

**Figure S4:**
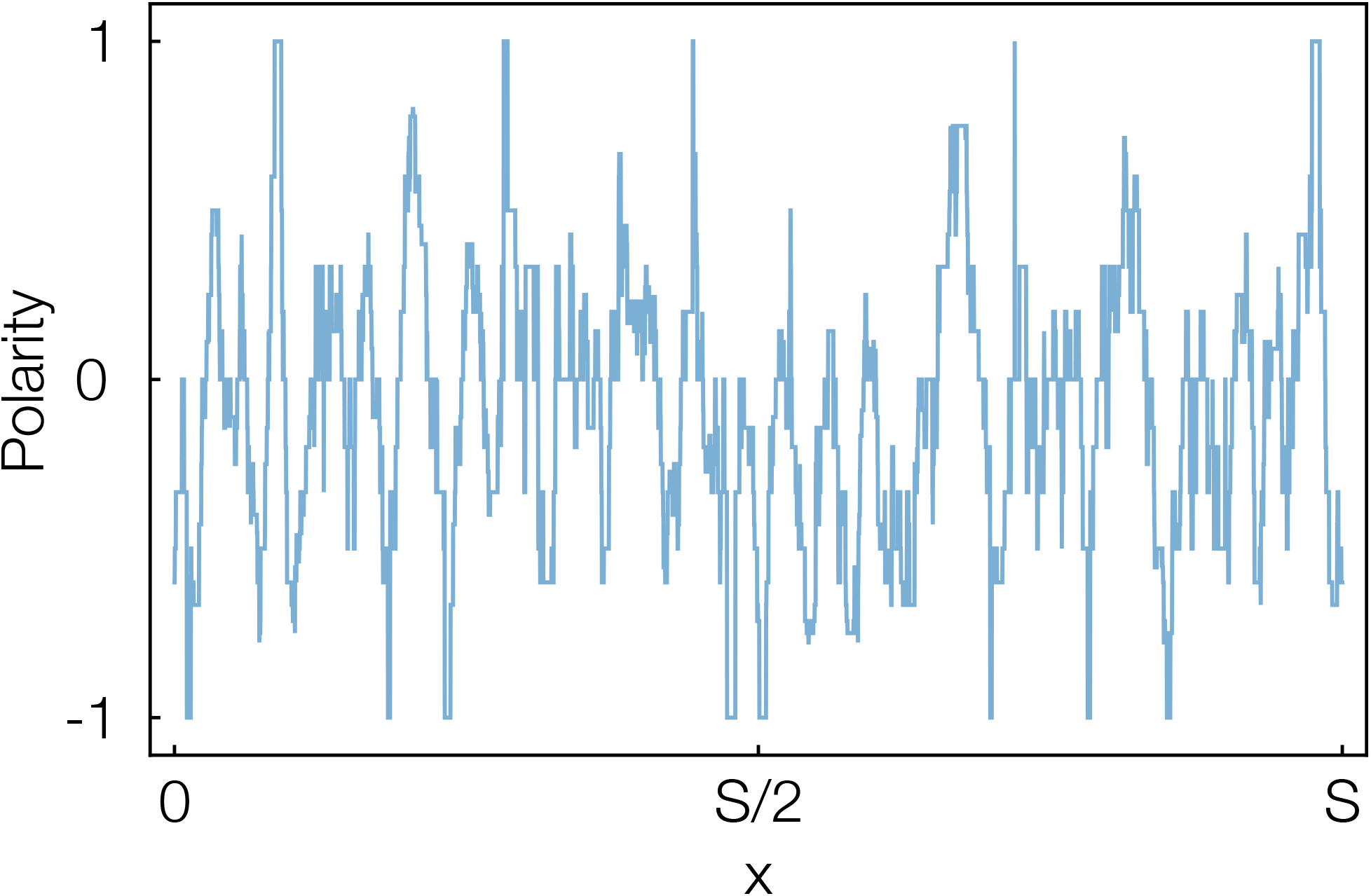
Exemplary polarity profile for a system of size *S* = 400μm with microtubules of length *L* = 6μm. The polarity profile is not completely random with independent values in all bins. Instead the polarities at different positions are correlated over a typical distance *L* (Eq. S34).

This equation implies that 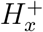 and 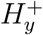 are independent if |*x* − *y*| ≥ *L* (then their covariance is zero), and that for |*x* − *y*| < *L* their correlation decays linearly with the distance. Taken together,

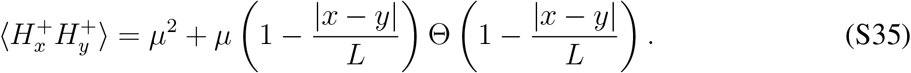

Due to symmetry, the same is true for 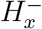. Furthermore, 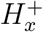 and 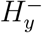 are independent for all *x*, *y*.

Equipped with these properties of the number of microtubules passing through the different positions 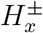, we now consider the local polarity. In terms of 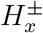 it is defined as

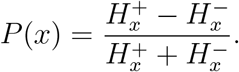

Due to the symmetry between left- and right-oriented microtubules, the average is

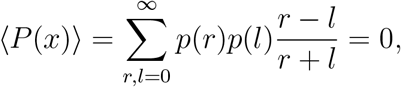

where we defined 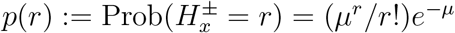.

The calculation of the other moments is a bit more involved, and we will repeatedly use the following identity for *a* ≥ 1 and *i* ∈ ℕ

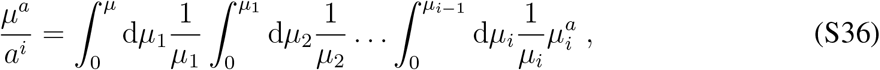

which is straightforwardly proven by induction and explicit integration of the right-hand side. Using this identity S36, the variance of the local polarity is given by

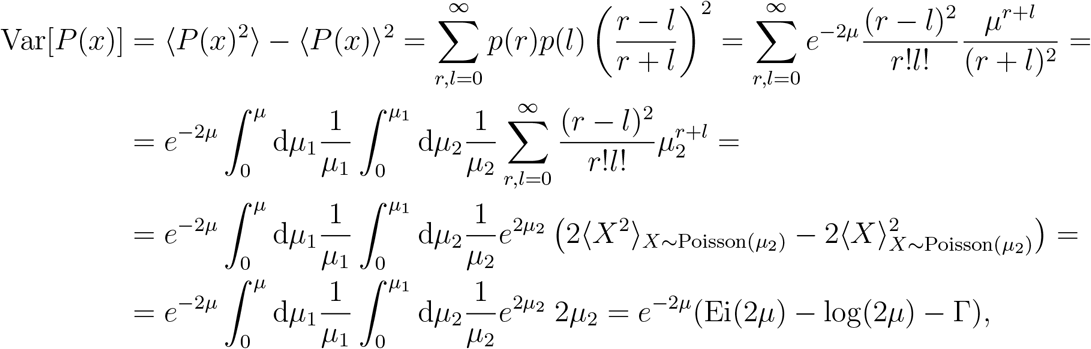

with the exponential integral Ei(x) and the Euler-Mascheroni constant Γ. Furthermore 〈*f*(*X*)〉_*X*~Poisson(*μ*)_ denotes the average of *f*(*X*) where *X* is distributed according to a Poisson distribution with parameter *μ*. In this calculation we used that 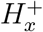 and 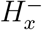 and, thus, the probabilities for *r* and *l*, *p*(*r*) and *p*(*l*), are independent.

In summary, we have

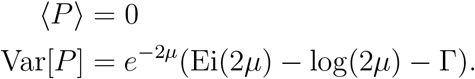

As expected, since the number of both types of microtubules (to the left and right, respectively) is on average the same everywhere, the mean of the polarity is zero. The variance of the local polarity Var[*P*] is monotonically decreasing for *μ* ≥ 1 and decays to 0 for large average densities *μ*, implying that the distribution of local polarities is less broad for larger values of *μ*. This result is intuitive as for higher *μ* more microtubules contribute to the local polarity and so the variance is smaller.

## Autocorrelation of the local polarity

Although the local polarity is already well characterized by its mean and variance (see Fig. 5A), for the distribution of the ambient polarity, information on the correlation between the local polarities at different positions is necessary. Thus, as a next step, we determine the autocorrelation of the local polarity

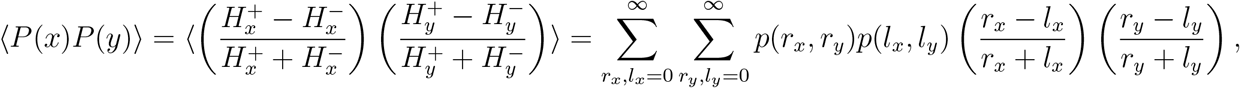

where 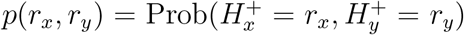 is the joint probability that 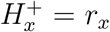 and 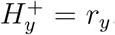. The random variables 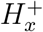 and 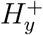 are independent only if |*x* − *y*| ≥ *L*, so the joint probability *p*(*r*_*x*_, *r*_*y*_) can be factorized as *p*(*r*_*x*_)*p*(*r*_*y*_) only in this case. Then, due to the symmetry between 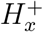 and 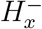 (and analogously for *y*), we have 〈*P*(*x*)*P*(*y*)〉 = 0.

If, however, |*x* − *y*| < *L*, such a factorization is not possible as 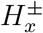 and 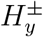 are correlated. To circumvent this difficulty, we split the random variables in two parts, namely one that describes the overlap/correlation of both, 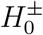, and one that captures the independent contributions, 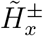 and 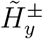:

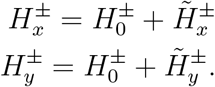

More concretely, we choose

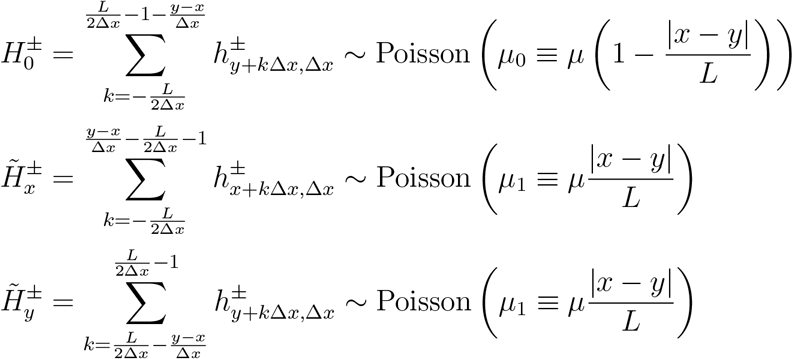

where we assumed *x* < *y* without loss of generality. The advantage of this decomposition is that now all 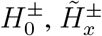 and 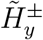 are independent (due to the independence of 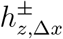 for different *z*).

For |*x* − *y*| < *L*, the autocorrelation of the local polarity is thus given by

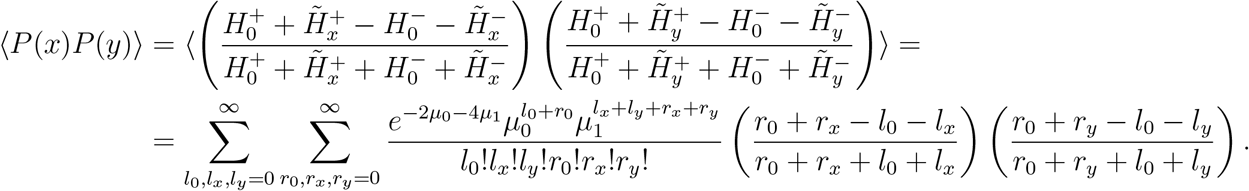

As we will see, we can make use of identity S36 again. In order to do so, we rewrite this expression in a slightly more complicated form

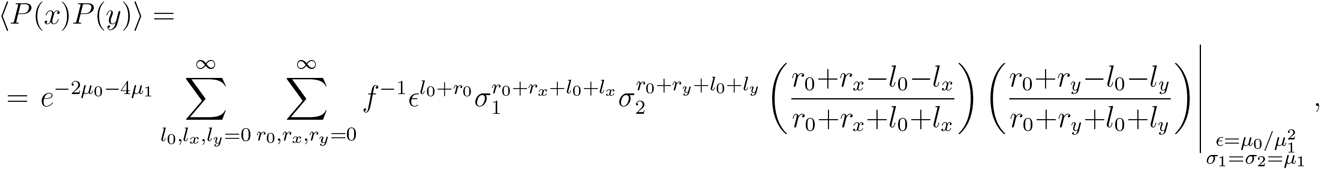

where we defined *f* = *f*(*l*_*i*_, *r*_*i*_) = *l*_0_!*l*_*x*_!*l*_*y*_!*r*_0_!*r*_*x*_!*r*_*y*_! as the product of the factorials. Applying now identity S36, we end up with

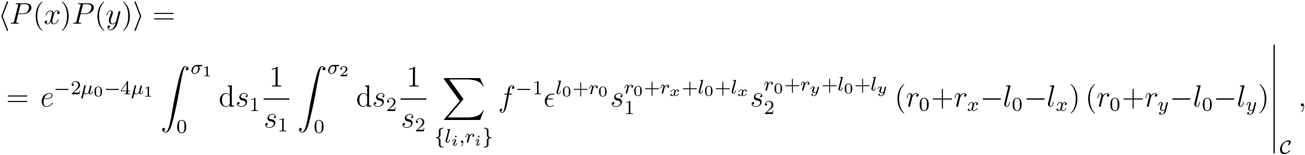

where the sum is over all *l*_*i*_, *r*_*i*_, *i* ∈ {0, *x*, *y*} from 0 to *∞* and 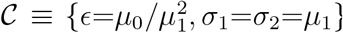. To continue, we rewrite the integrand as follows

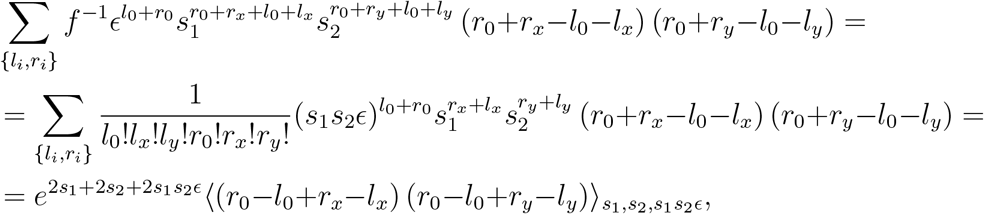

where the average 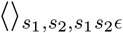 has to be interpreted with respect to the six independent variables *l*_*x*_, *r*_*x*_ ~ Poisson(*s*_1_), *l*_*y*_, *r*_*y*_ ~ Poisson(*s*_2_) and *l*_0_, *r*_0_ ~ Poisson(*s*_1_*s*_2_*ϵ*). Due to the independence and as 〈*r*_*x*_−*l*_*x*_)〉=〈*r*_*x*_−*l*_*x*_〉=0 the average is given by

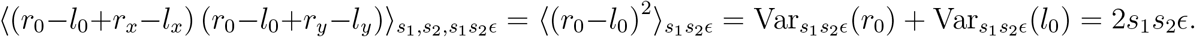

Taken together, we determine the autocorrelation of the local polarity to be

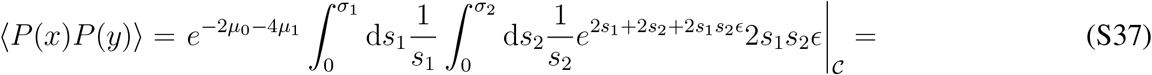

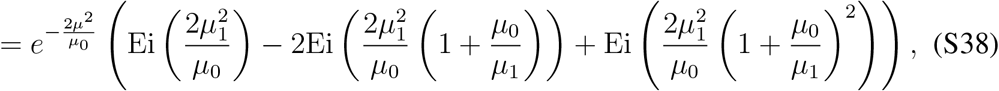

where we used that *μ*_0_ + *μ*_1_ = *μ*. Ei(*x*) denotes the exponential integral as before. When plotting the autocorrelation 〈*P*(*x*)*P*(*y*)〉 against the normalized distance |*x* − *y*|/*L* (using that *μ*_0_ = *μ*(1 − |*x* − *y*|/*L*) and *μ*_1_ = *μ*|*x* − *y*|/*L*), one realizes that the autocorrelation decays approximately linearly with the distance: 〈*P*(*x*)*P*(*y*)〉 ~ 1 − |*x* − *y*|/*L*. The origin of the approximately linear decay is that the number of microtubules directly linking two points decreases linearly with the distance as well. The proportionality constant for the linear decay, which corresponds to the limit for |*x* − *y*| → 0 (*μ*_0_ → *μ*, *μ*_1_ → 0), is calculated to be

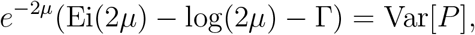

where we used that Ei(*x*) ≈ Γ + log(*x*) for small *x*. This is consistent with the result for the variance of the local polarity: lim_|*x*−*y*|→0_〈*P*(*x*)*P*(*y*)〉 = Var[*P*].

In summary, we thus find for the autocorrelation of the local polarity

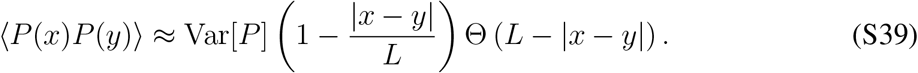

This equality implies that the local polarities at positions less than one microtubule length apart are not independent. Instead, the finite length of microtubules introduces correlations as one microtubules contributes to the polarity at different locations. The strength of the correlations decays approximately linearly with the distance, analogously to the number of microtubules directly connecting the two locations. In the following, we will use this result to determine the distribution of the ambient polarity.

## Distribution of the ambient polarity

Analogously to the local polarity, we characterize the distribution of the ambient polarity by its average and variance, implicitly assuming that the ambient polarity is reasonably well described by a normal distribution. In the calculations, however we do not explicitly use that the distribution is approximated by a normal distribution and our results are valid irrespective of this assumption.

From the definition of the ambient polarity 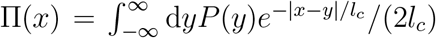, its average and variance are given by

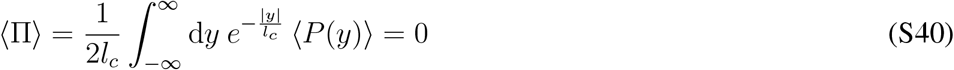

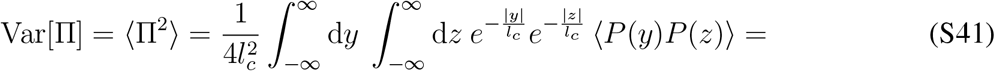

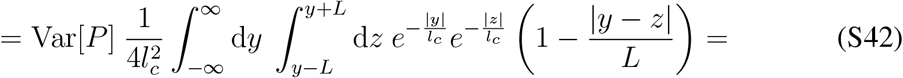

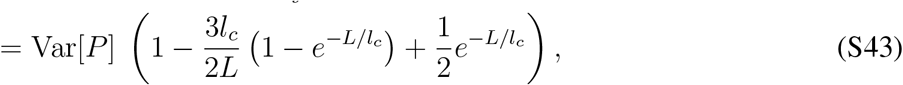

where - due to the translational symmetry - we considered *x* = 0 without loss of generality. From Eq. 5a in the main text, *v*^(±)^/*V*_*m*_ = ±(1 − *α*)(1 ∓ Π), we immediately conclude that the variance of the normalized velocity *v*^(±)^/*V*_*m*_, Var[*v/V*_*m*_], is given by

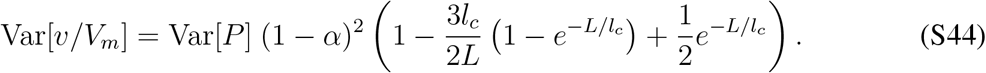

Taking the limit *l*_*c*_ → 0, we find Var[Π] = Var[*P*] = Var[*v/V*_*m*_]/(1 − α), and, conversely, for *l*_*c*_ → ∞, we have Var[Π], Var[*v/V*_*m*_] → 0. These limits illustrate our intuition that for small characteristic length only the local polarity matters, whereas for large characteristic length all microtubules feel the same ambient polarity.

## S6 Stochastic, agent-based simulation of the *in silico* experiment

For the *in silico* experiment in the main text, a random polarity profile was generated as described in Appendix “*In silico* experiment: Random polarity field”. The velocities are then determined from Eqs. 5a, 5b according to the continuum description. In this continuum approximation, all microtubules at one position exhibit exactly the same speed. This assumption will not be satisfied in experimental filament gels where two microtubules at the same location by chance can be connected to a different set of microtubules and thus experience a different environment. As a result, not only microtubules at different locations show a different speed but also the speed of microtubules at the same location can vary, leading to a broader distribution of the microtubule velocities. The goal of this Appendix is to gauge the strength of this effect.

To this end, the *in silico* experiment is performed again in terms of a stochastic, agent-based simulation for the same parameters as in the main text (Fig. 5): Compared to the *in silico* experiment in the main text, not only the microtubules are randomly distributed but also the interactions between microtubules are randomly chosen. That is, each microtubule randomly interacts with on average *N* of its neighbors, and the individual velocities are determined from the force-balance equations 3. Figure S6A shows the measured probability distribution for all combinations of local polarity and velocity, analogously to Fig. 5. As in the main text, the histograms for both quantities were obtained as projections of the density plot to the respective axis. Since in the agent-based simulations not all microtubules at one position *x* exhibit the same velocity, here, “local velocity” refers to the average velocity of all equally-oriented microtubules passing through position *x*. As expected, the distribution of the average velocity in the stochastic *in silico* experiment is broader than the velocity distribution in the main text which agreed very well with the prediction from our theory (black lines). The ratio of both standard deviations is approximately *σ*[*v/V*_*m*_]/*σ*[*P*] = 0.28, indicating that the stochastic nature of the interactions indeed influences the filament velocities.

**Figure S5:**
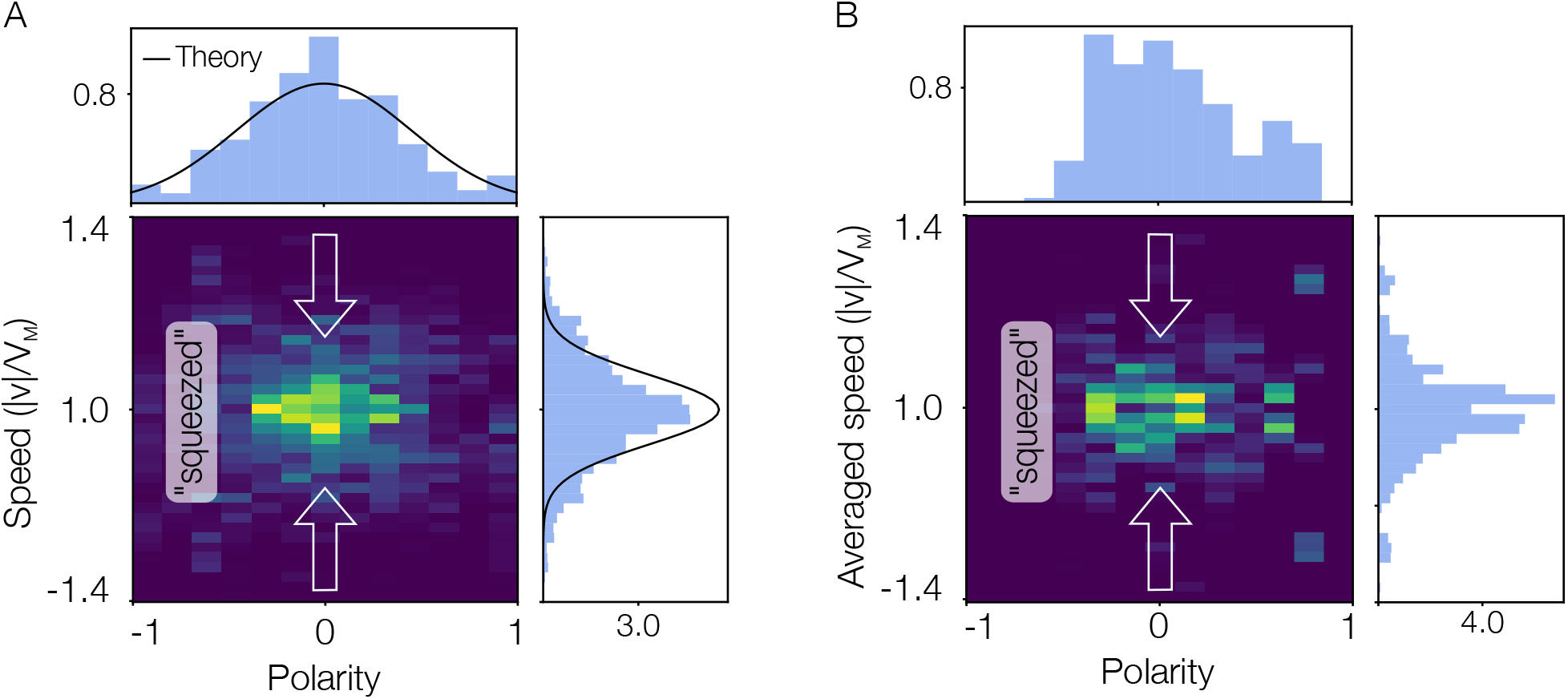
Density plot and histograms for the local polarity and speed analogous to Fig. 5 but measured in the stochastic agent-based simulation. Measurements are taken for 6000 independent filament gels. The parameters are identical to the ones used in the main text. (**A**) The local speed |*v*^(±)^| is calculated as the average speed of all equally-oriented microtubules passing through the respective position. Compared to the prediction from the nonlocal continuum theory (black lines), the variance of the velocity is slightly larger. (**B**) The local averaged speed |*v*^(±)^| is calculated as the average speed of all equally-oriented microtubules that are part of a region of 4μm in the center of the filament gel. Compared to the measurement in (**A**), the variance of both the polarity and the velocity is smaller.

In experiments with filament gels, average velocities at single points *x* in space usually can not be unravelled. Instead, one measures the average polarity and velocity of regions of the order of several μm. This averaging is expected to lead to a narrower distribution of polarities and velocities as compared to measurements of the polarities and velocities of single points in space. To estimate the influence of this averaging procedure, we perform a second measurement in the stochastic *in silico* experiment. The goal of this second version is to mimic a typical measurement in experiments, so we do not record the average velocities of single points in space. Instead, we measure the average velocity and polarity of all equally oriented microtubules that are part of a region of 4μm in the center of the filament gel of length 400μm. Performing this measurement for 6000 independent filament gels yields a distribution of polarities and velocities as shown in the density plots and histograms in Fig. S6B. As can be seen from a comparison of Figs. S6A and B, the distributions for the polarity and velocity after averaging over a range of several μm is noticeably smaller. Indeed, the result of the average measurements in the stochastic, agent-based *in silico* experiment is similar to the one shown in the main text. This finding implies that the expected broadening and narrowing of the velocity distribution due to stochasticity and averaging, respectively, more or less balance. Of course, this depends on choices such as the size of the region the velocity and polarity are averaged over. In the *in silico* experiment we chose it to be 4μm, similar to the width of the photo-bleached region in *in vitro* filament gels ^18^. Generally, the width of the velocity distribution is expected to depend on experimental details but we anticipate that independent of these details the width of the velocity distribution decreases with increasing characteristic length.

## S7 Extension of the *in silico* experiment to a broader class of systems

The analysis in section “The agent-based model can describe the weak velocity-polarity sensitivity” in the main text, and the corresponding construction of the polarity field in the *in silico* experiment, Appendix “*In silico* experiment: Random polarity field”, are based on the assumption that the polarity field does not have any spatial structure. That is, the system is translationally invariant and -on average - all positions are equivalent. However, generally, this premise will not be fulfilled. In this appendix, we thus want to extend our previous analysis to a broader class of systems.

## Class of systems

As we have seen before in Appendix “*In silico* experiment: Random polarity field”, the variance of the ambient polarity, Var[Π], depends on the autocorrelation of the local polarity 〈*P*(*x*)*P*(*y*)〉. Thus, in order to make any statements about the distribution of the ambient polarity, we need to make some assumptions on the correlation of the local polarities at different locations. The most obvious property of the system that leads to correlations of the local polarities at different locations is the finite extension of microtubules. As discussed before, due to the finite microtubule length *L* > 0, an excess of microtubules at one position leads to an excess of microtubules at distances maximally *L* apart. In the following, we will assume that this contribution to the correlation dominates, and that there are only weak correlation effects, for instance due to filament dynamics or feedback. We believe that in this case it is reasonable to assume that the covariance of the local polarity at different positions (the autocorrelation) decays linearly with distance up to |*x* − *y*| = *L*:

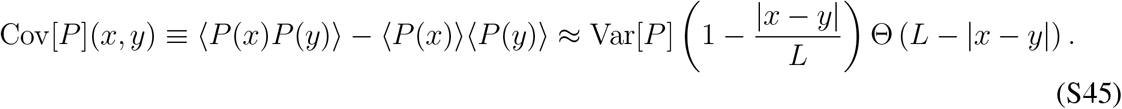

Here we furthermore assumed that the magnitude of the fluctuations in the polarity is similar everywhere: Var[*P*] is approximately spatially invariant.

We will restrict our discussion to this class of system as quantitative statements for the general case are difficult to obtain.

## Prediction

Let us consider systems where Eq. S45 holds. Suppose we measure the spatial profile of the average of the local polarities and of the velocities, 〈*P*(*x*)〉 and 〈*v*^(±)^(*x*)〉, where the average denotes an ensemble average at fixed position *x*. Moreover, we determine the average variance of the local polarity Var[*P*] = 〈Var[*P*(*x*)]〉_*x*_, where Var[*P*(*x*)] = 〈*P*(*x*)^2^〉 − 〈*P*(*x*)〉^2^ is the variance of the local polarity at fixed position *x* and 〈〉_*x*_ denotes an average over all locations *x*. Then, our theory predicts that the covariance of the velocity at different positions is

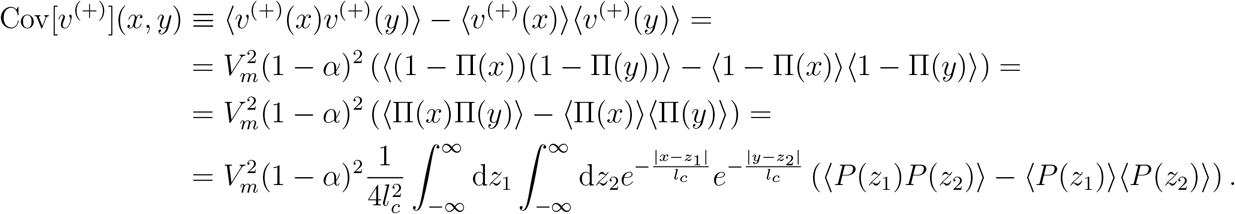

Using assumption S45, this expression becomes

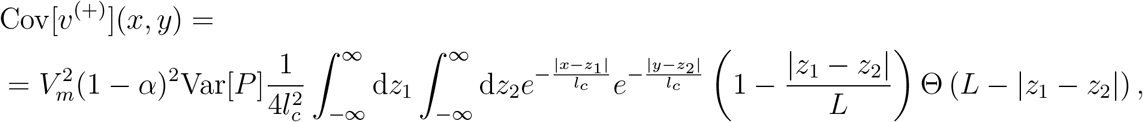

which holds for the velocities *v*^(−)^ as well. For general distance *x* − *y*, the analytic expression is not very insightful and the expression is best understood graphically.

Fig. S6 shows a comparison between the normalized covariance of the local polarities (the autocorrelation coefficient), Cov[*P*](*x, y*)/Var_*P*_, and of the velocity, Cov[*v*^(±)^](*x*, *y*)/Var[*v*^(±)^], for different *l*_*c*_. Whereas the correlation of the local polarity quickly decays to zero (after a distance |*x* − *y*| = *L*), the correlation of the velocities is much more long-ranged and its correlation length increases with *l*_*c*_.

The covariance of the velocity for *x* = *y*, Cov[*v*^(+)^](*x*, *x*), which corresponds to the variance of the velocity Var[*v*^(±)^], is given as

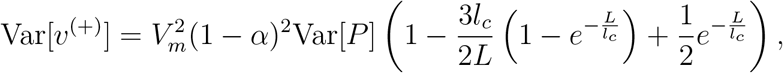

in terms of the variance of the local polarity, Var[*P*]. Similarly, the variance of the ambient polarity, Var[Π], is given by

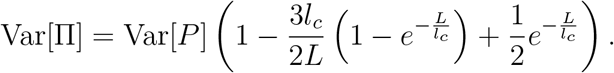

So, for the broader class of systems considered here we recover exactly the same result as for the *in silico* experiment, Eq. S43.

**Figure S6:**
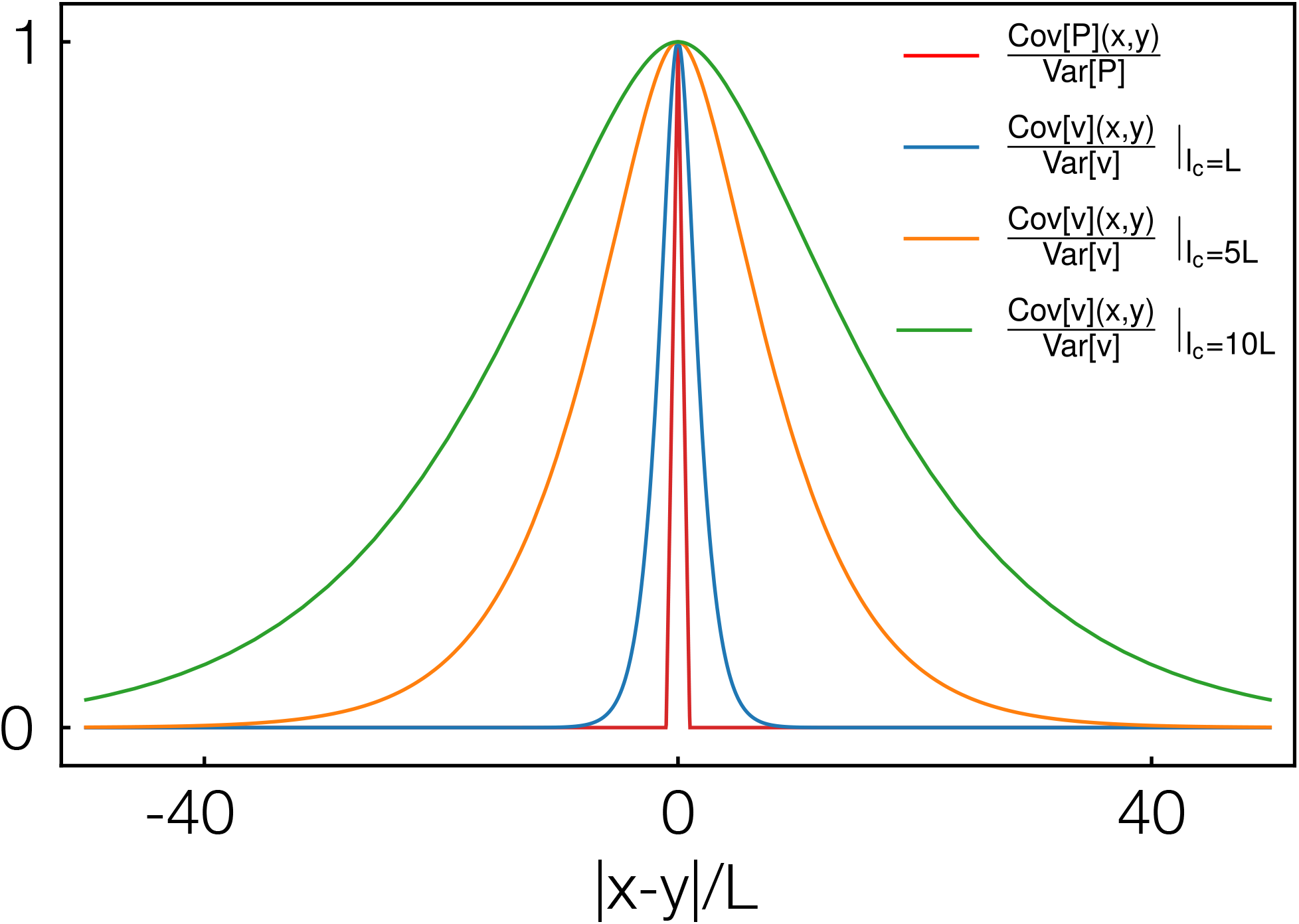
Comparison of the autocorrelation coefficient of the local polarity, Cov[*P*](*x*, *y*)/Var[*P*], and of the velocity, Cov[*v*^(±)^](*x*, *y*)/Var[*v*^(±)^], for different values of the characteristic length *l*_*c*_. The velocity correlations decay much slower as compared to the correlations in the local polarity. The correlation length of the velocity scales with *l*_*c*_, that is, for larger *l*_*c*_, the correlations length is larger as well.

## S8 Comparison of our results for small characteristic length to the dilute limit

One of the central results of our work is that there is an intrinsic length scale of the system that determines the velocity-polarity relation. This characteristic length *l*_*c*_ captures how far information on the local forces propagates through the network. Naively, one can argue that for dilute systems *l*_*c*_ is small and, correspondingly, that forces only act locally. This conclusion fits well with the intuitive conception of a dilute limit where filaments are arranged in disconnected patches. Nonetheless, one has to be careful to directly compare our result to the dilute limit. With regard to this limit, there are two main assumptions in our continuum theory.

1. *Single patch:* All microtubules are directly or indirectly (via other microtubules) connected to each other and there are no disconnected patches of microtubules. This assumption corresponds to hypothesizing that the filament network works above the percolation threshold and that the average number of interaction partners *N* is not too small.
2. *Sufficient interaction partners:* The number of neighbors a microtubule interacts with is limited by the average number of interaction partners *N* and not by the number of neighbors. This assumption is based on the idea that there is always a sufficient number of neighbors (possible interaction partners) present for each microtubule. Instead of linearly depending on the microtubule densities, the force thus exhibits a dependency on the fraction *φ*^(±)^ (Eq. 3).

Both assumptions do not necessarily apply to the dilute limit. We believe that the second assumption regarding a sufficient number of interaction partners can be readily relaxed also within a continuum description. For this purpose, one could, for instance, try to incorporate a phenomenological term *Nρ*^(±)^/(*N* + *ρ*^+^ + *ρ*^−^) for the number of interactions with (±) microtubules. Such a term would converge to *Nφ*^(±)^ for large total density *ρ* ≡ *ρ*^+^ + *ρ*^−^ ≫ *N*, as used in our description. Conversely, for small *ρ* it captures a linear dependency of the number of (±) interactions on the respective density. Taken together, investigating how such an effective term changes the behavior should be instructive for a more quantitative understanding of the dilute limit.

The first assumption is conceptually more difficult to overcome within a continuum description. But indeed, for parameters estimated as in section “The agent-based model can describe the weak velocity-polarity sensitivity”, regions with zero density (and thus disconnected patches) occur regularly in our stochastic, agent-based simulation. These empty spaces arise due to the stochastic loss of connections between microtubules and a following drifting apart of different patches. Vice versa, such empty spaces stochastically vanish again if two patches meet. The interplay of these opposing, stochastic processes leads to patch boundaries that are not static but change randomly.

We suppose that one could effectively incorporate this behavior into our continuum theory. To this end, one might first consider systems of finite sizes and then try to average their behavior with regard to exponentially distributed system sizes. Intuitively, we would expect that this procedure leads to an enhanced effective attenuation and thus to a lower effective value of the characteristic length *l*_*c*_ but does not change our results qualitatively: Due to the stochastic loss of connections between patches (particularly for systems close to the percolation threshold), there is not only loss of information due to drag but also abruptly at the boundaries of the patches. If these boundaries are fluctuating, the abrupt loss at the boundaries is on average smoothed and should be qualitatively comparable to a continuous loss of information by drag.

Overall, we think that - despite these assumptions of our theory - our results help bridging the gap between previous findings for dilute and heavily crosslinked filament networks.

## Footnotes

1 for zero overall polarity

2 Care has to be taken when comparing our result to the dilute limit: We restricted our discussion to the case of sufficient number of interaction partners, whereas usually for dilute systems disconnected patches of filaments are considered. For a more comprehensive discussion on how our results are related to results for dilute systems we refer the reader to the appendix “Comparison of our results for small characteristic length to the dilute limit”.

3 Note that we ignore sets of measure zero.

